# Bioinformatics and next generation sequencing data analysis to identify key genes and pathways influencing in Parkinson’s disease

**DOI:** 10.1101/2022.02.27.482208

**Authors:** Basavaraj Vastrad, Chanabasayya Vastrad

## Abstract

Parkinson’s disease (PD) is the most commonly diagnosed neurodegenerative disorder Identification of novel prognostic and pathogenesis biomarkers plays a pivotal role in the management of the PD. Next generation sequencing (NGS) dataset from the GEO (Gene Expression Omnibus) database were used to identify differentially expressed genes (DEGs) in PD. Gene Ontology (GO) and REACTOME pathway enrichmental analyses were performed to elucidate the functional roles of the DEGs. Protein-protein interaction (PPI), modules, miRNA-hub gene regulatory network and TF-hub gene regulatory network were established. The receiver operating characteristic curve (ROC) analysis was used to explore the diagnostic values of hub genes in PD. In total, 957 DEGs were identified, of which 478 were up regulated genes and 479 were down regulated genes. GO and pathway enrichment analysis results revealed that the up regulated genes were mainly enriched in nervous system development, cell junction, transporter activity and neuronal system, whereas down regulated genes were mainly enriched in response to stimulus, cell periphery, identical protein binding and immune system. The top hub genes in the constructed PPI network, modules, miRNA-hub gene regulatory network and TF-hub gene regulatory network were OTUB1, PPP2R1A, AP2M1, PIN1, USP11, CDK2, IQGAP1, NEDD4, VIM and CDK1. Furthermore, ROC analysis showed that hub genes were having good diagnostic values. We identified a series of essential genes along with the pathways that were most closely related with PD initiation and progression. Our results provide a more detailed molecular mechanism for the advancement of PD, shedding light on the potential biomarkers and therapeutic targets.

## Introduction

Parkinson’s disease (PD) is the second most common type of neurodegenerative disorder and a leading cause of impairment of voluntary motor control, which places a great burden on the economy of health and reduces quality of life [1]. PD accounts for 1-2 per 1000 of the population at any time [2]. PD involves the degeneration of dopaminergic neurons in the substantia nigra of the midbrain and the advancement of neuronal Lewy Bodies [3]. Numerous factors might affect PD progression, including genetic factors [4], aging and inflammatory factors [5], and environmental factors [6]. However, how these factors affect the development of PD requires further investigation and no effective method has been developed for treatment and diagnosis. It is therefore urgent to identify novel diagnostic and prognostic biomarkers for PD.

Molecular biology investigations have identified numerous biomarkers and signaling pathways that contribute to PD, including the brain-derived neurotrophic factor (BDNF) [7], histone deacetylase 4 (HDAC4) [8], vacuolar protein sorting 35 (VPS35) [9], leucine-rich repeat kinase 2 (LRRK2) [10], phosphatidylinositol binding clathrin assembly protein (PICALM) [11], RhoA-ROCK signaling pathways [12], Nrf2 signaling pathways [13], GTPase-p38 MAPK signaling pathways [14], JAK/STAT signaling pathway [15] and PI3K/Akt signaling pathway [16]. Further investigation into the molecular events linked with PD is required.

With the advance of the human genome project, PD has been investigated at the genetic level. Next generation sequencing (NGS) technology can be used to find genes that cause early PD. NGS technology has the characteristics of high sensitivity. NGS technology is extensively used in disease diagnosis [17] and novel therapeutic target screening [18]. At present, NGS technology was used to find potential biomarkers that affect the advancement of diseases in studies [19]. NGS technology plays an important role in elucidating gene expression in PD [20]. However, the pathogenesis of PD remains unclear.

In this investigation, we explored novel biomarkers for PD diagnosis and targeting therapies. We manipulated the NGS data of GSE135036 [21] dataset from the GEO (Gene Expression Omnibus) (http://www.ncbi.nlm.nih.gov/geo/) [22] database to distinguish differentially expressed genes (DEGs) between PD and normal control. gene ontology (GO) and pathway enrichment analysis were done to elucidate the functions of the DEGs. The hub genes, miRNAs (micro RNA) and TFs (transcription factors) related to the pathogenesis of PD were chosen by protein-protein interaction (PPI) network, modules, miRNA-hub gene regulatory network and TF-hub gene regulatory network. Receiver operating characteristic curve (ROC) analysis was performed to validate the hub genes, which could be used as molecular biomarkers or diagnostic or therapeutic targets for PD therapy. Collectively, our investigation will help the advancement of a genetic diagnosis for PD and more effective measures of prevention and interference.

## Materials and methods

### Data resources

The GEO database is a public genome database. In this investigation, NGS dataset GSE135036 [21] was downloaded from the GEO database. GSE135036 was based on Illumina NextSeq 500 (Homo sapiens) platform. GSE135036 dataset contained 25 samples, including 13 PD samples and 12 normal control samples.

### Identification of DEGs

The DESeq2 package of R software [23] was used for the screening of differentially expressed genes (DEGs) with the criteria of fold change > 0.513 for up regulated genes, fold change < −0.61 for down regulated genes and adjusted P < 0.05. The results were visualized as a volcano plot and heat map using the ggplot2 and gplot in R software.

### GO and pathway enrichment analyses of DEGs

We used GO analysis (http://www.geneontology.org) [24] to explore the potential functions of the DEGs. GO is a widely used bioinformatics tool to identify genes and study-related biological processes. Three terms comprised the GO analysis, including cellular component (CC), biological process (BP), and molecular function (MF). We employed KEGG to search for potential pathways of the overlapping DEGs. REACTOME (https://reactome.org/) [25] is a database to study gene functions and pathways from big datasets sourced from high-throughput experiments. g:Profiler (http://biit.cs.ut.ee/gprofiler/) [26], an online biological database, was used to analyze the GO and REACTOME terms. A value of p < 0.05 was considered significant.

### Construction of the PPI network and module analysis

The HIPPIE interactome (http://cbdm-01.zdv.uni-mainz.de/~mschaefer/hippie/index.php) is a database for searching between known proteins and predicting the interactions between proteins [27]. We used it to build PPI network for DEGs. The interaction networks were visualized with Cytoscape software version 3.8.2 (http://www.cytoscape.org/) [28]. Centrality analysis includes analyzing the degree [29], betweenness [30], stress [31] and closeness [32] of network nodes. Cytoscape plug-in Network Analyzer was used to calculate the values of degree, betweenness, stress and closeness to predict the key genes [33]. Functional modules in the network were identified by using the plug-in PEWCC1 [34] of Cytoscape.

### miRNA-hub gene regulatory network construction

miRNet database (https://www.mirnet.ca/) [35] is a comprehensive database, which provides the largest available set of predicted and experimentally validated miRNA-hub gene interactions. Additionally, it not only records miRNA binding sites in the entire sequence of genes but also compared this information with the binding sites of 14 existing miRNA-hub gene prediction programs: TarBase, miRTarBase, miRecords, miRanda (S mansoni only), miR2Disease, HMDD, PhenomiR, SM2miR, PharmacomiR, EpimiR, starBase, TransmiR, ADmiRE, and TAM 2.0. Cytoscape software version 3.8.2 [28] was used to construct a miRNA-hub gene regulatory network and analyze the interactions of the miRNAs and hub genes.

### TF-hub gene regulatory network construction

NetworkAnalyst database (https://www.networkanalyst.ca/) [36] is a comprehensive database, which provides the largest available set of predicted and experimentally validated TF-hub gene interactions. Additionally, it not only records TF binding sites in the entire sequence of genes but also compared this information with the binding sites of existing TF-hub gene prediction program Jasper. Cytoscape software version 3.8.2 [28] was used to construct a TF-hub gene regulatory network and analyze the interactions of the TFs and hub genes.

### Receiver operating characteristic curve (ROC) analysis

Receiver operating characteristic (ROC) curves were adopted to analyze the diagnostic value of the hub genes for PD. To check hub genes’ diagnostic values, we plotted ROC curves and determined area under the curve (AUC) with “pROC” R package [37]. The hub genes with the highest AUC value were consistent as having the key power for diagnosing PD.

## Results

### Identification of DEGs

To explore the role of systems biology in the pathogenesis of PD, we analyzed NGS data of GSE135036 by DESeq2 package of R software. NGS results from GSE135036, candidate DEGs were screened using the criteria of fold change > 0.513 for up regulated genes, fold change < −0.61 for down regulated genes and adjusted P < 0.05. There were 957 DEGs in GSE135036. These DEGs included 478 up regulated genes and 479 down regulated genes between PD and normal control samples (Fig. 1 and Table 1). Hierarchical clustering analysis revealed a clear distinction of DEGs between patients with PD and normal control (Fig. 2).

**Fig. 1.**
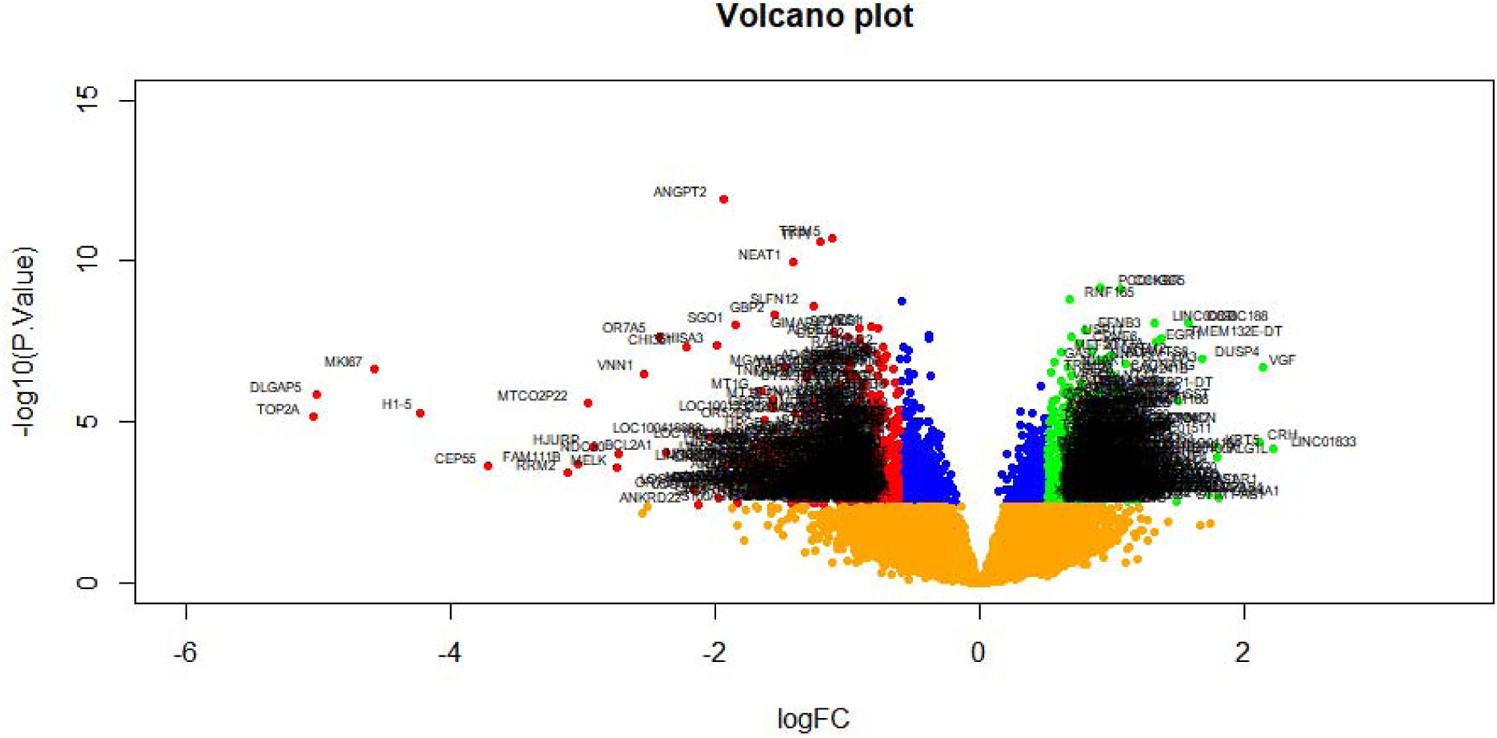
Volcano plot of differentially expressed genes. Genes with a significant change of more than two-fold were selected. Green dot represented up regulated significant genes and red dot represented down regulated significant genes.

**Fig. 2.**
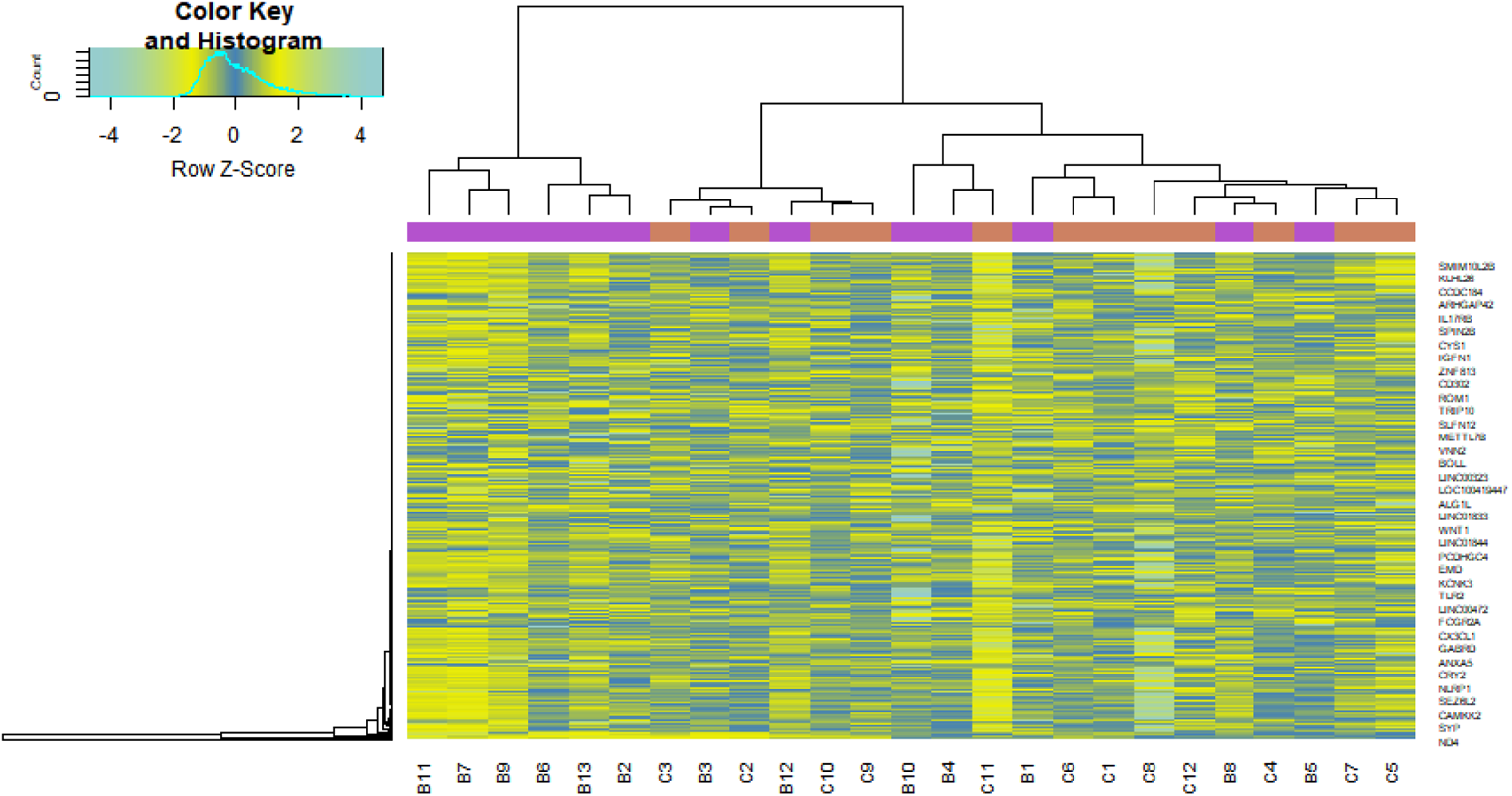
Heat map of differentially expressed genes. Legend on the top left indicate log fold change of genes. (A1 – A37 = normal control samples; B1 – B47 = HF samples)

**Table 1.**
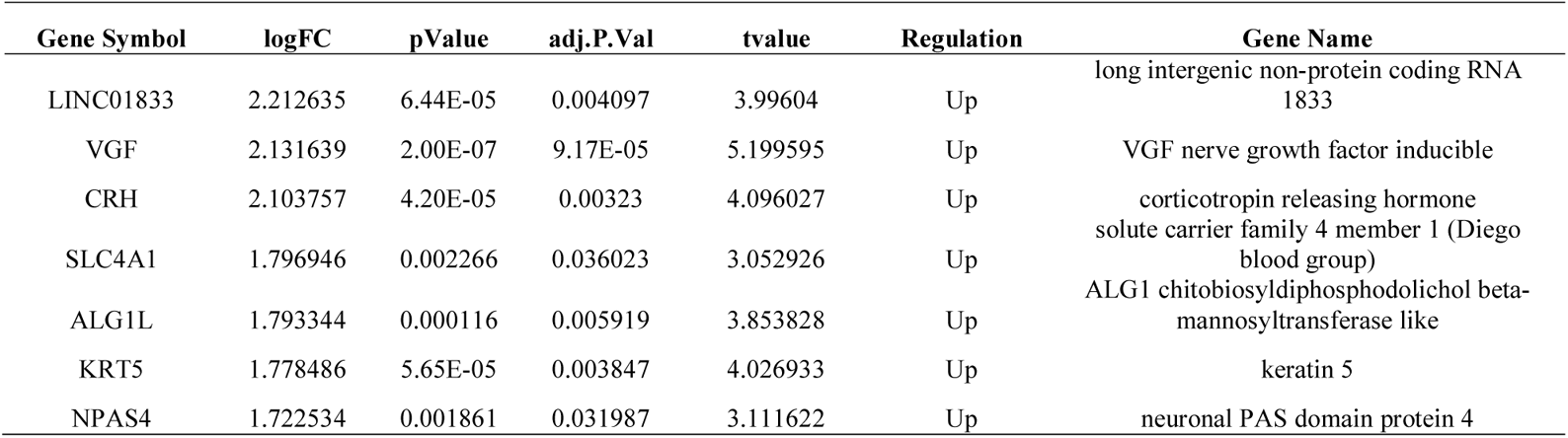
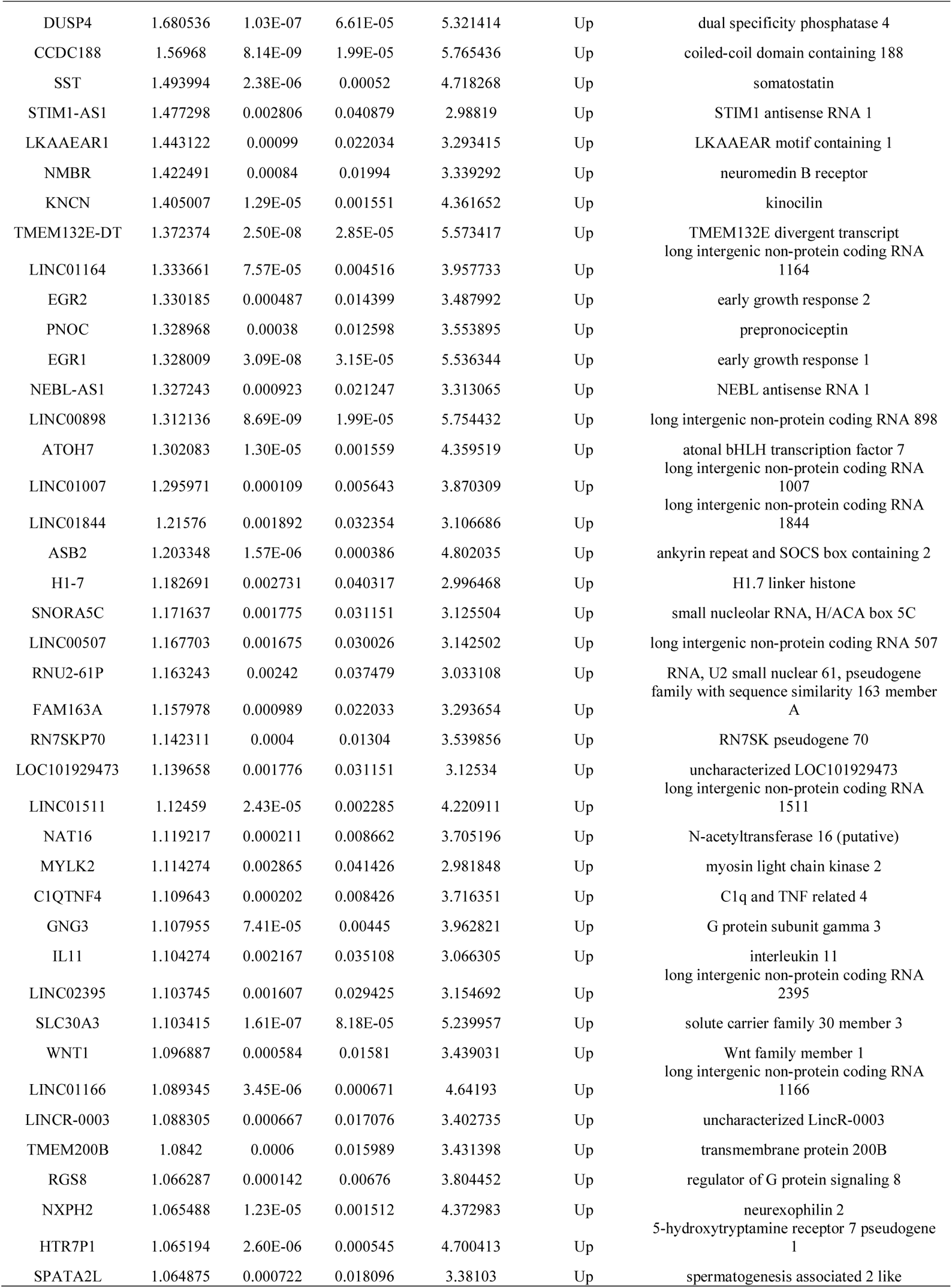
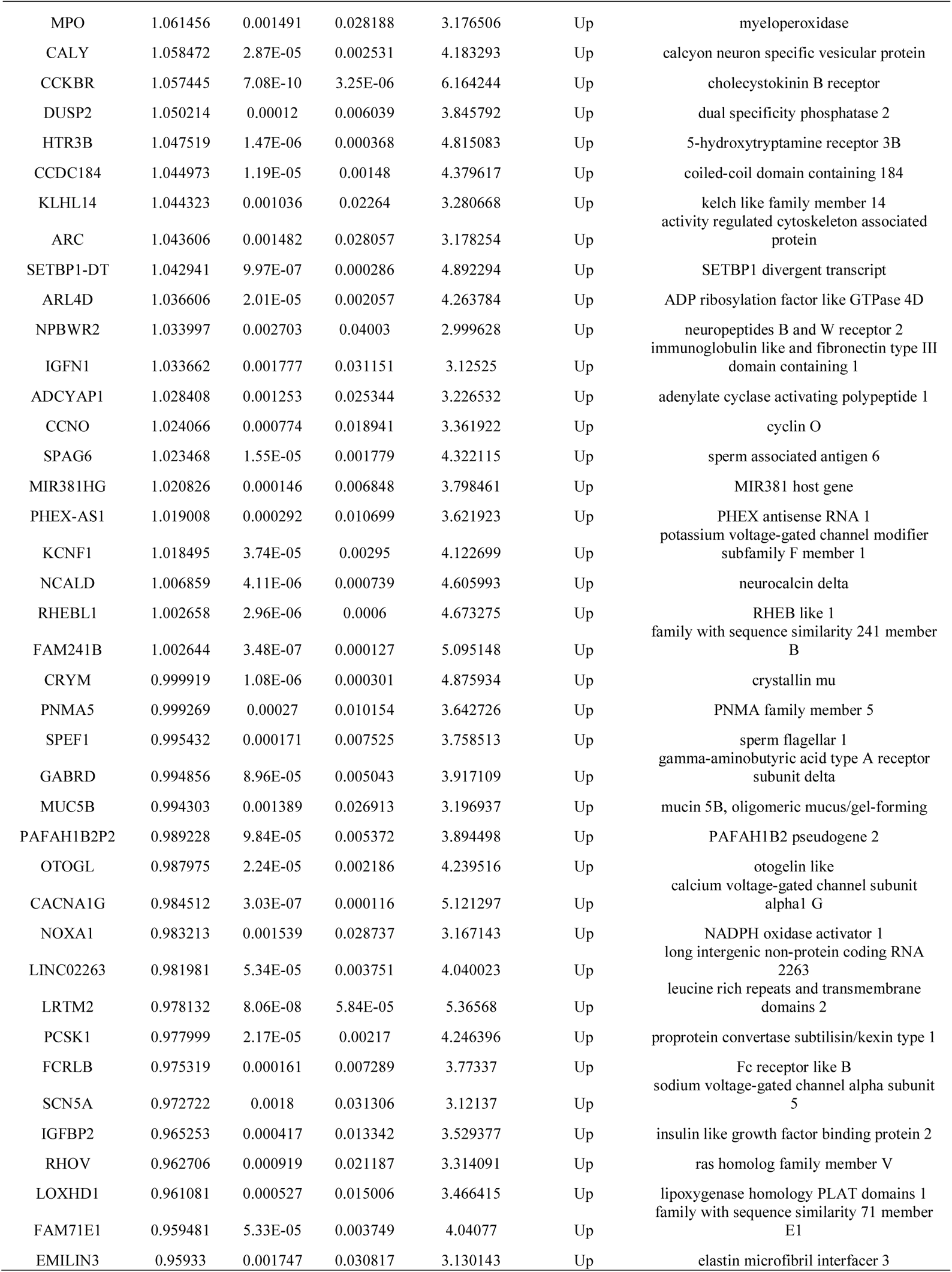
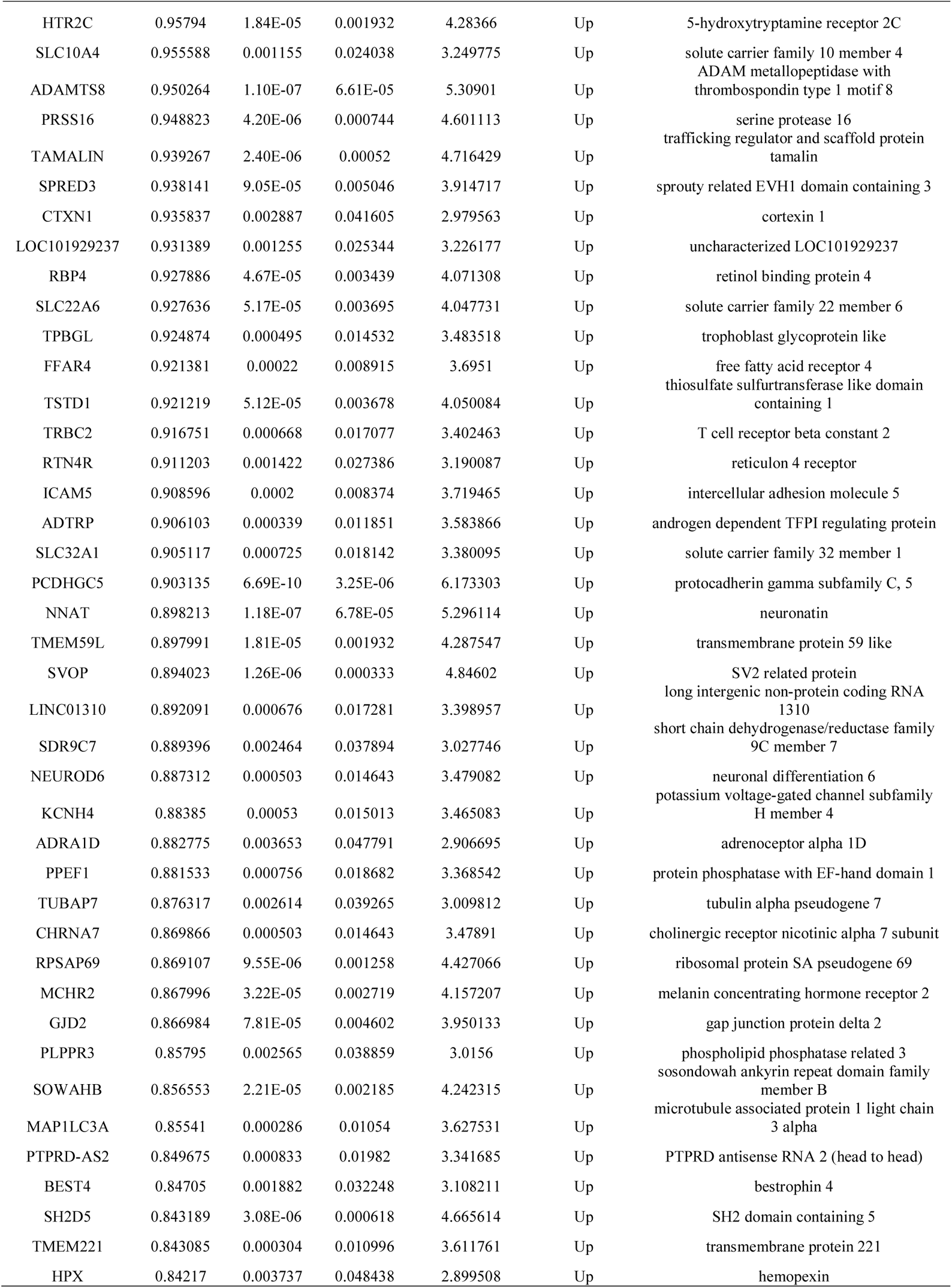
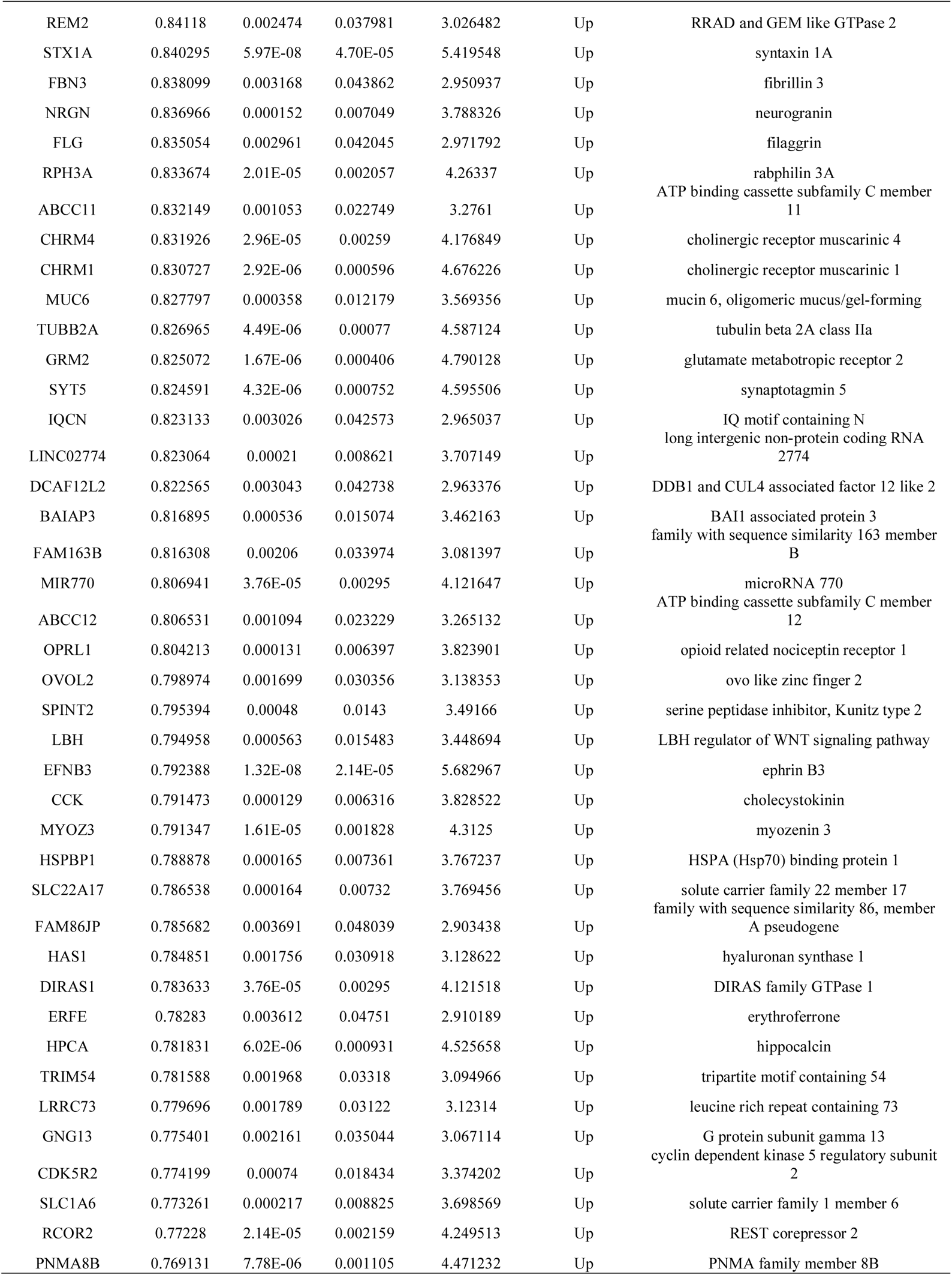
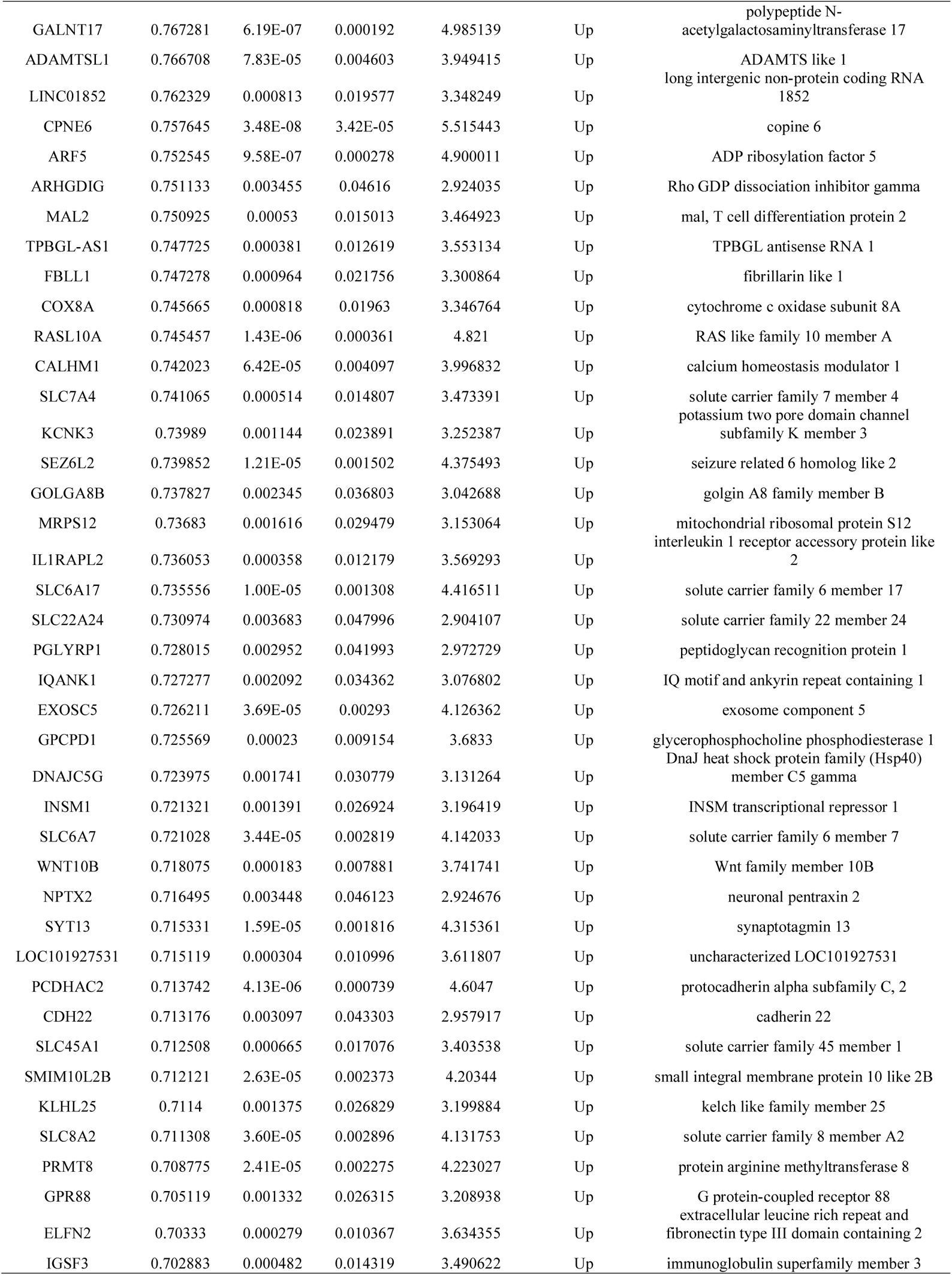
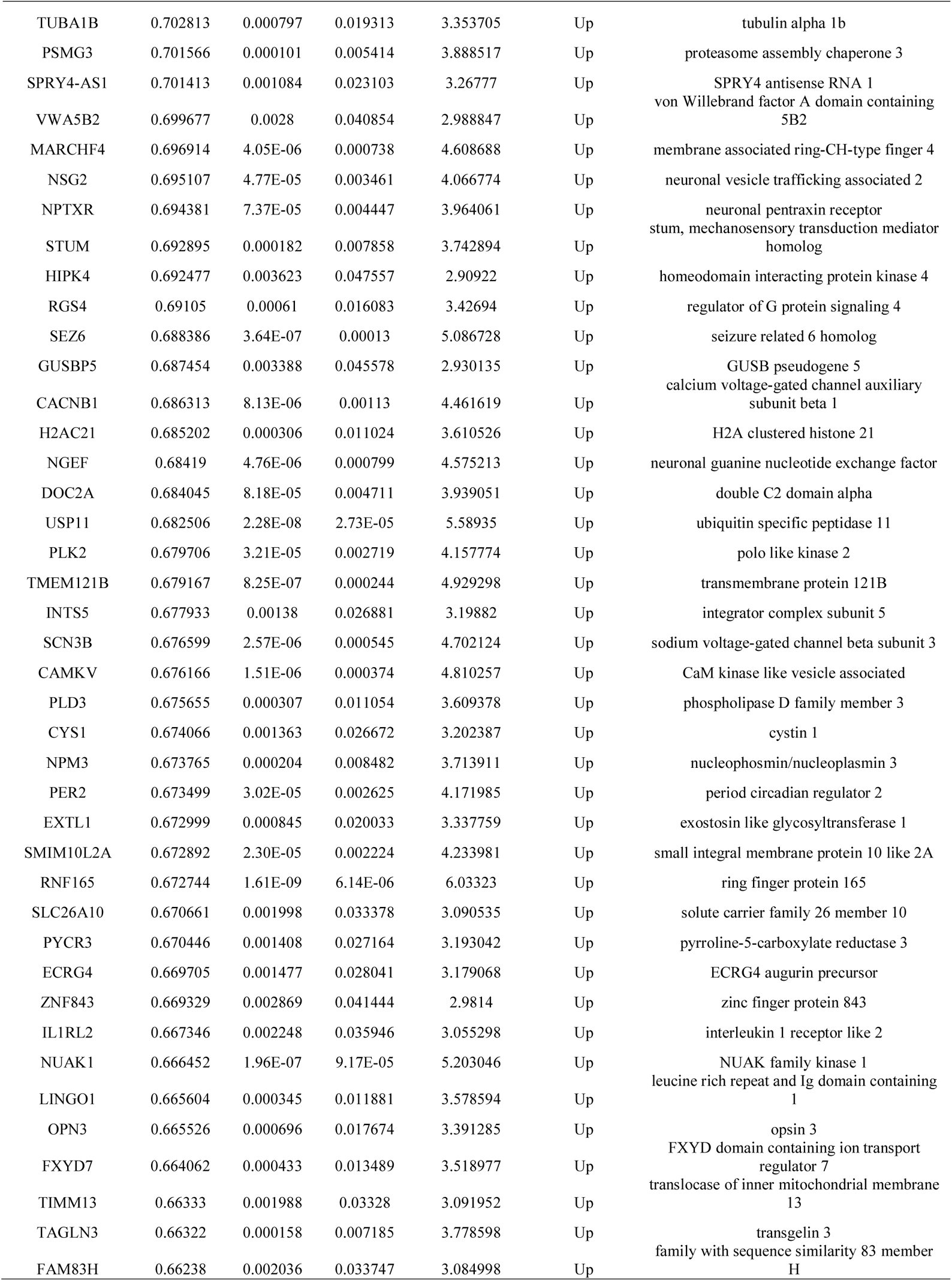
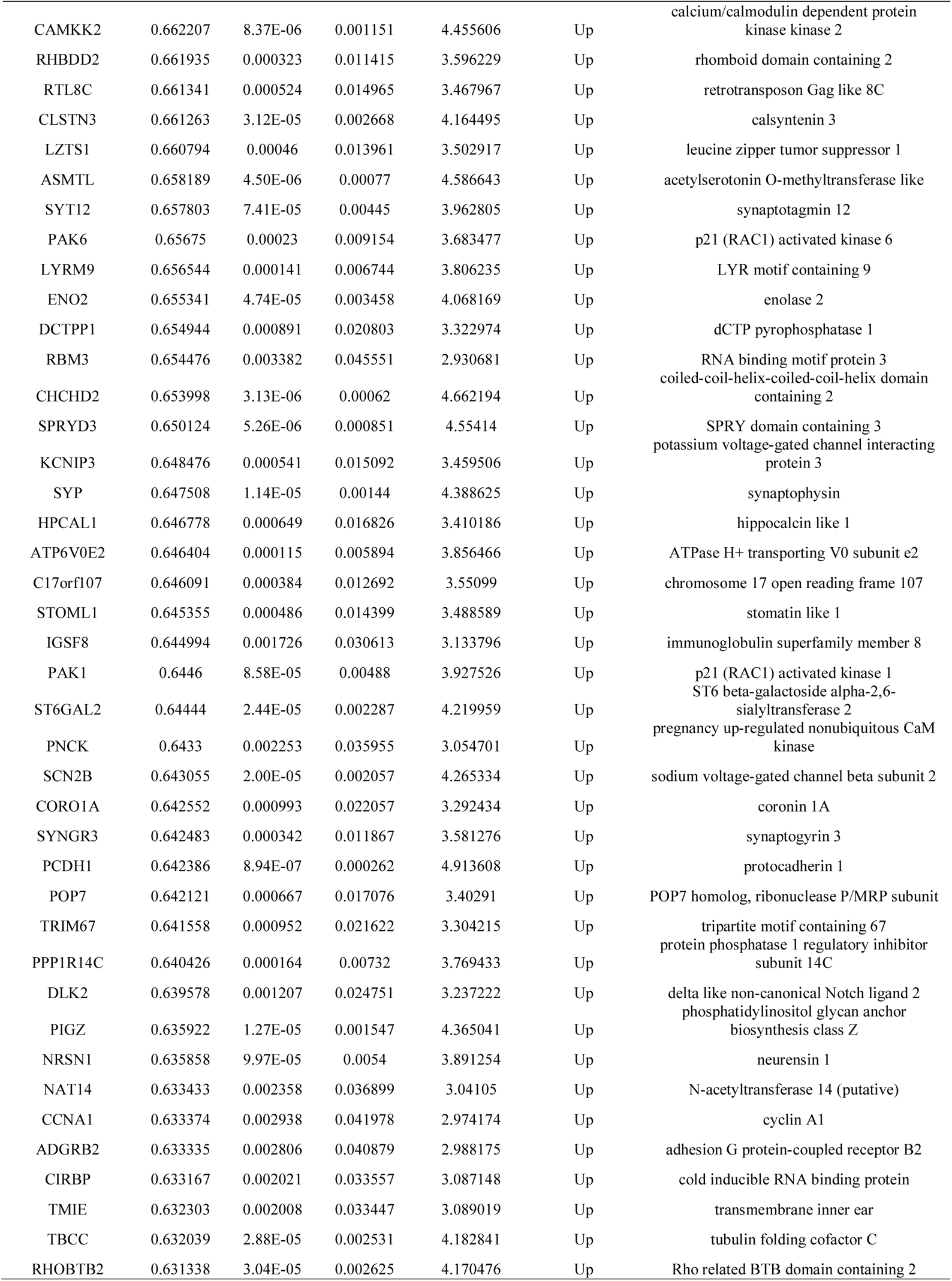
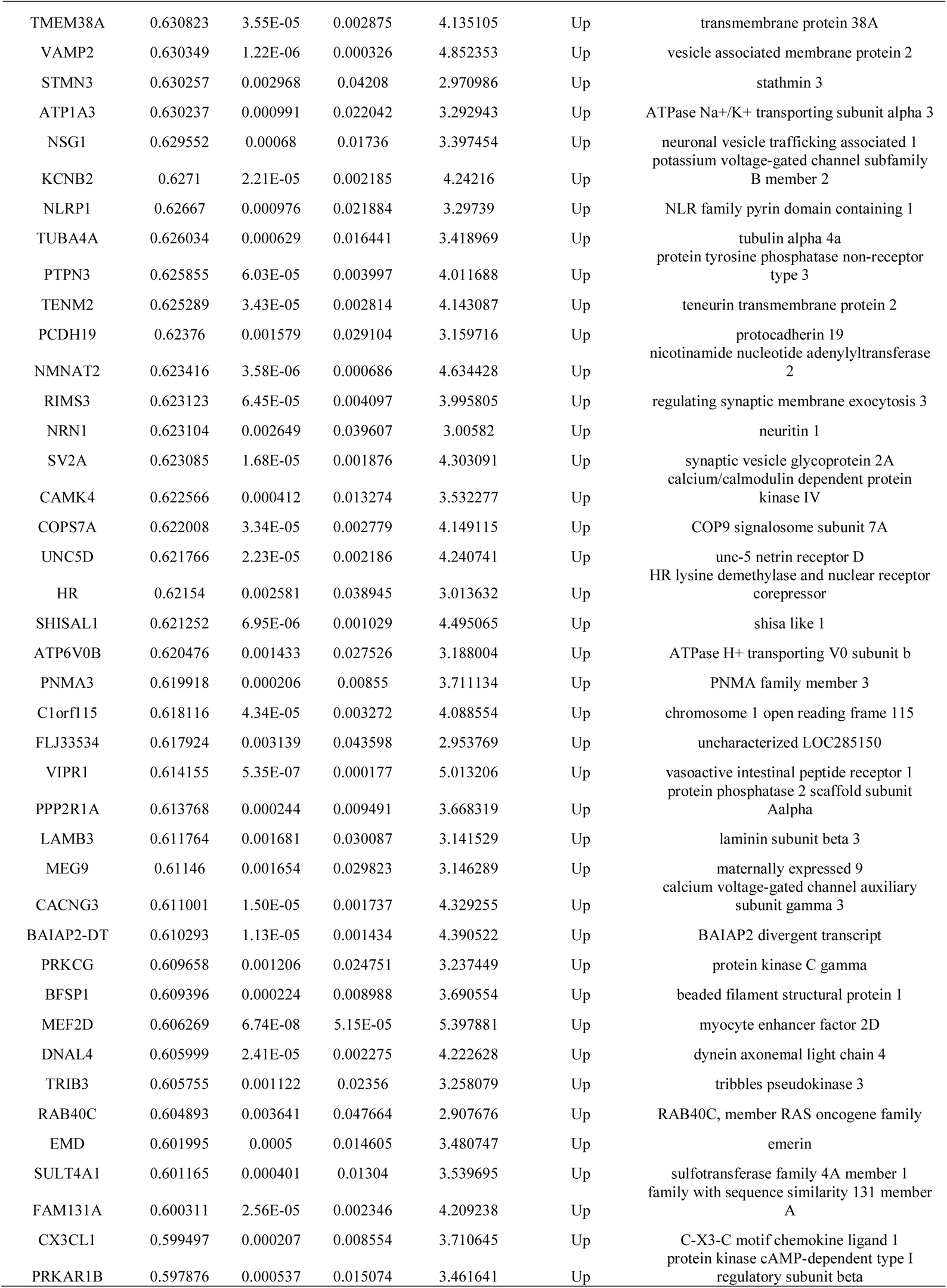
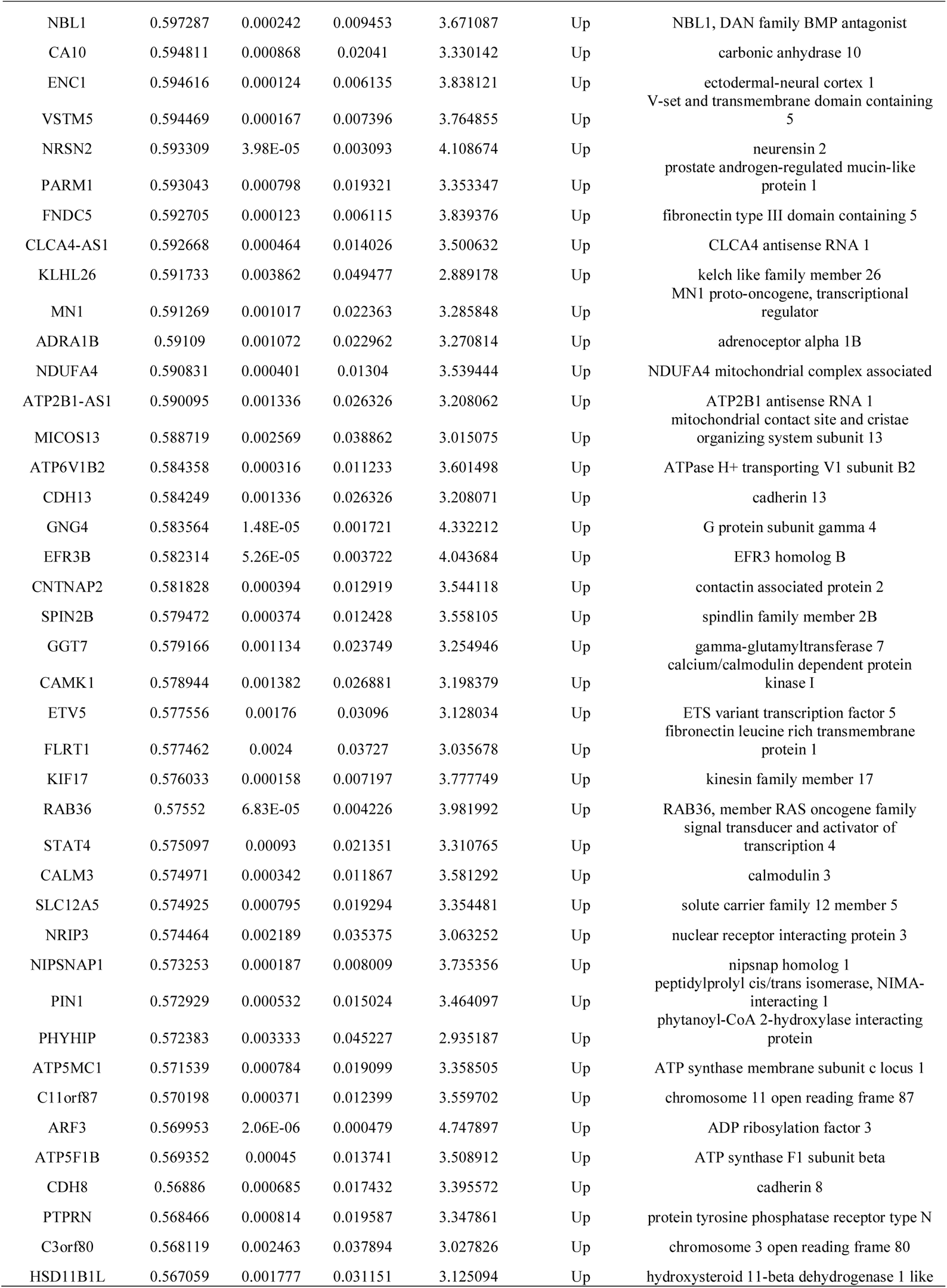
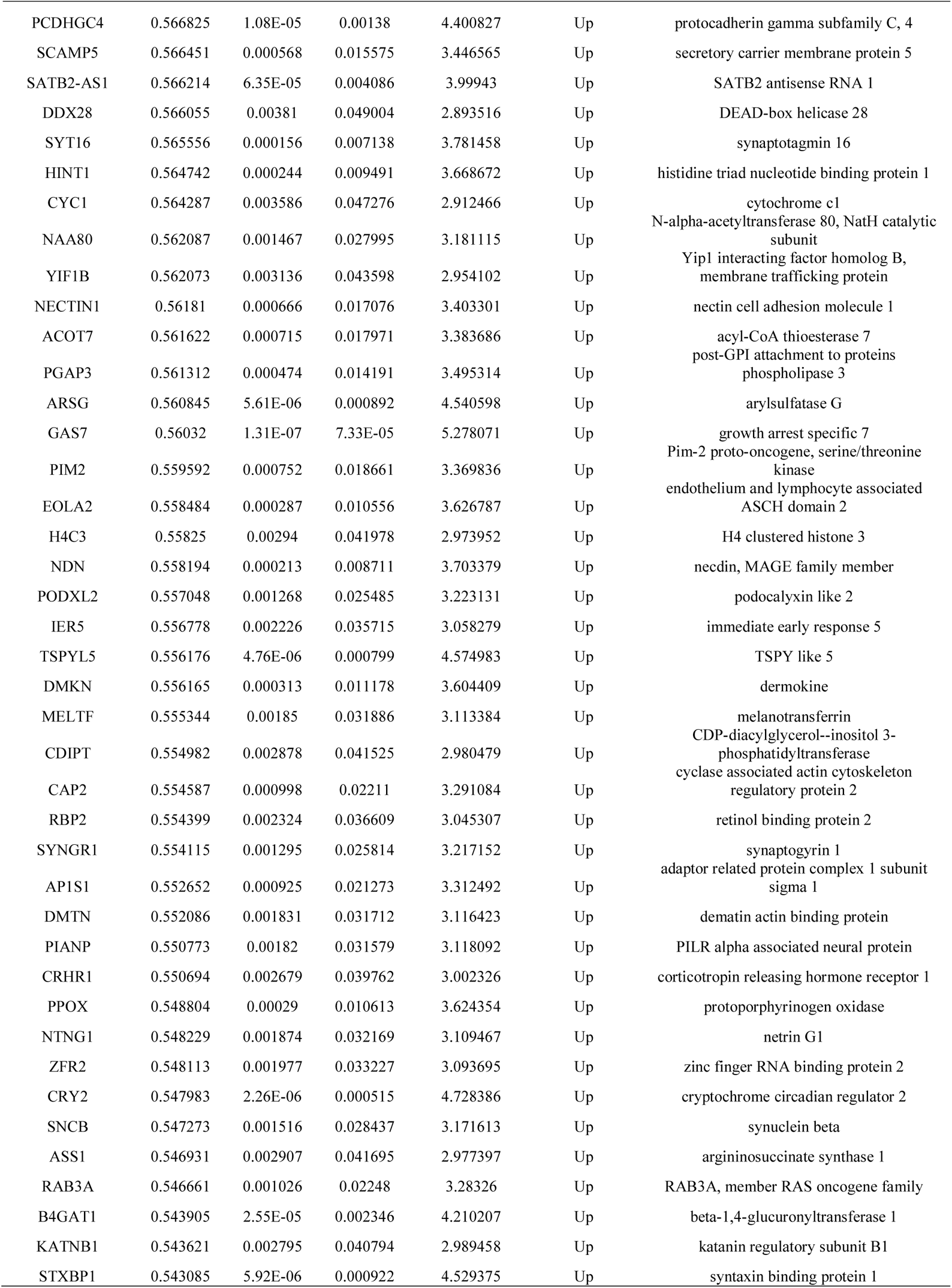
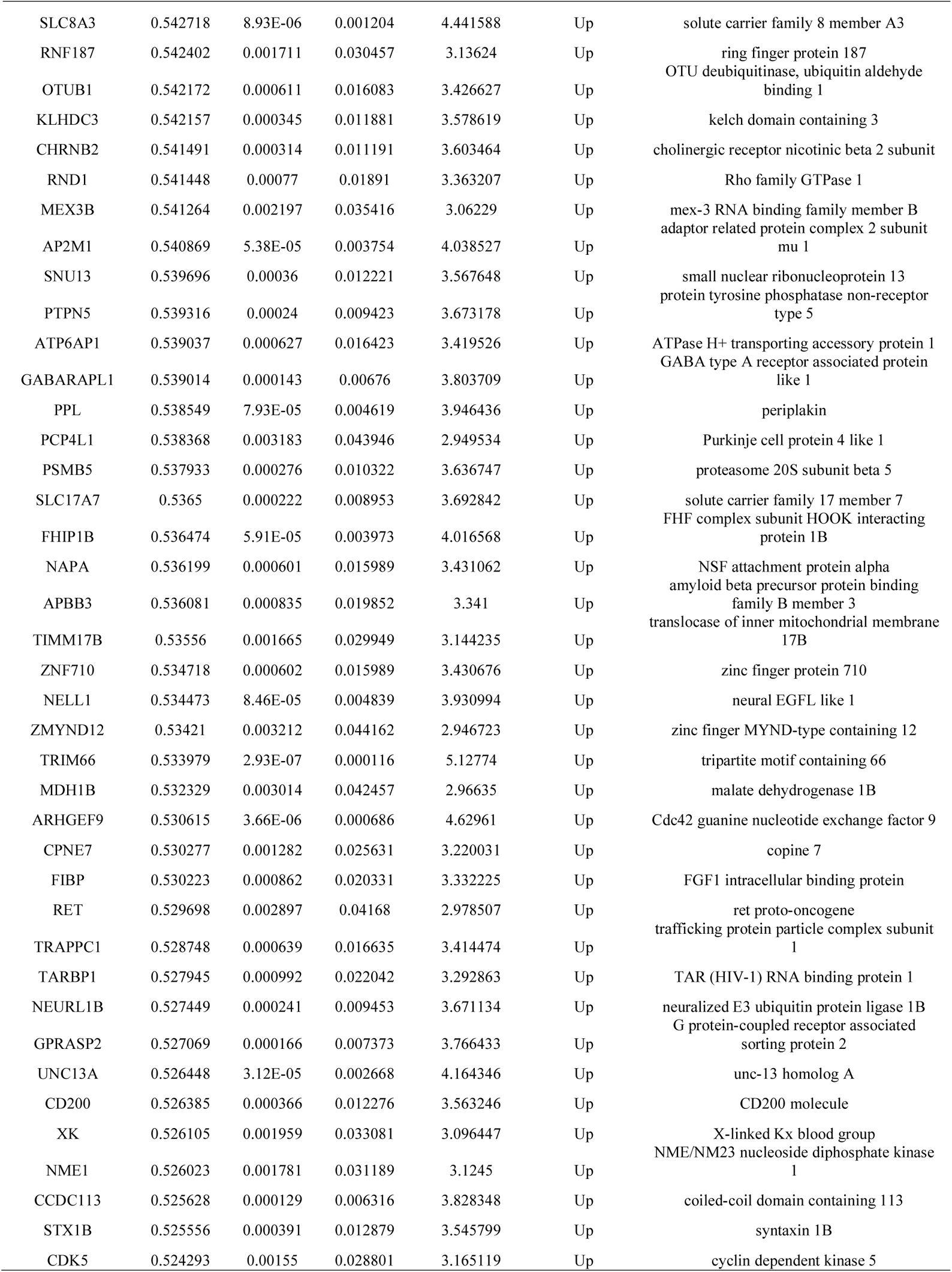
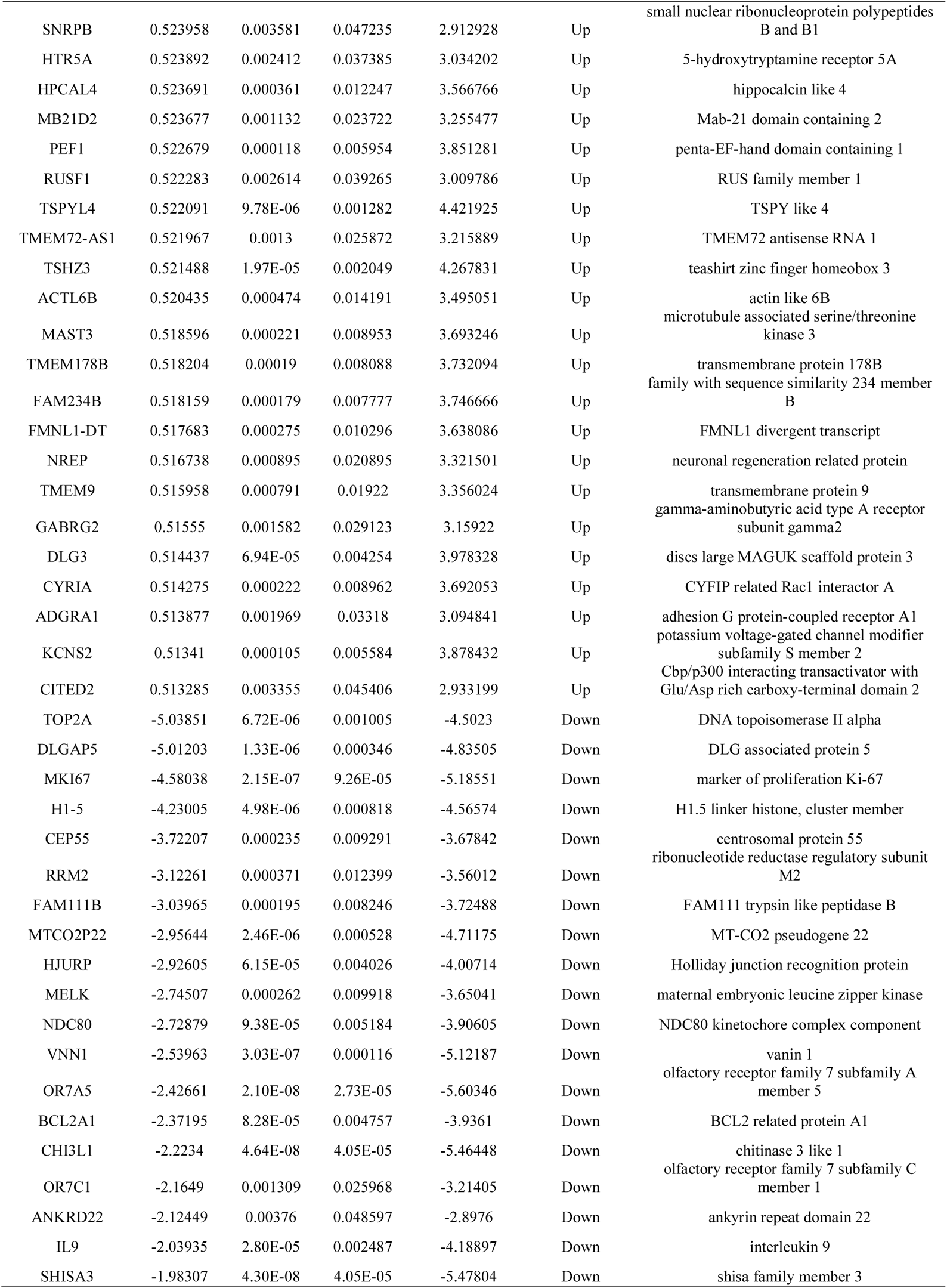
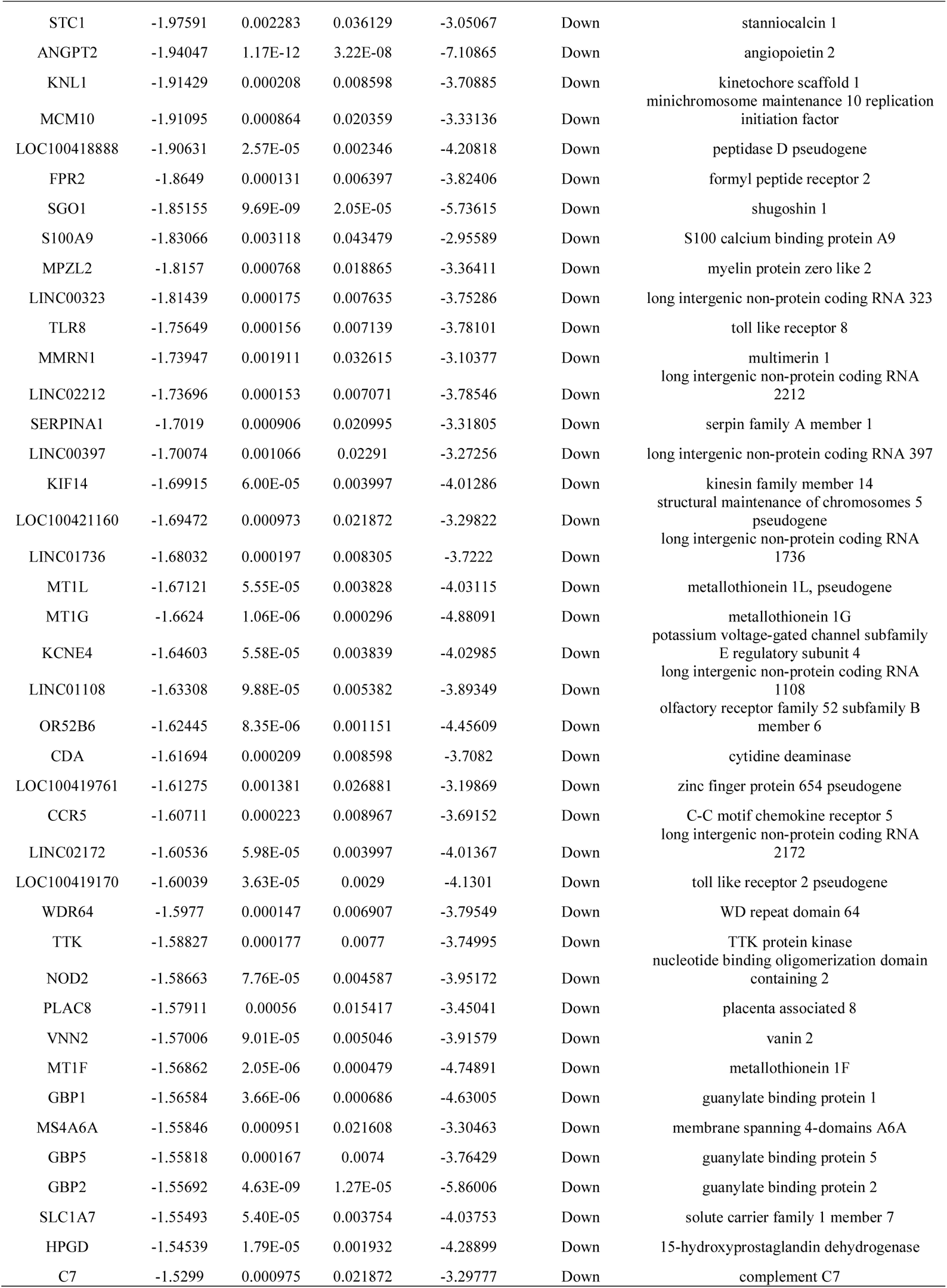
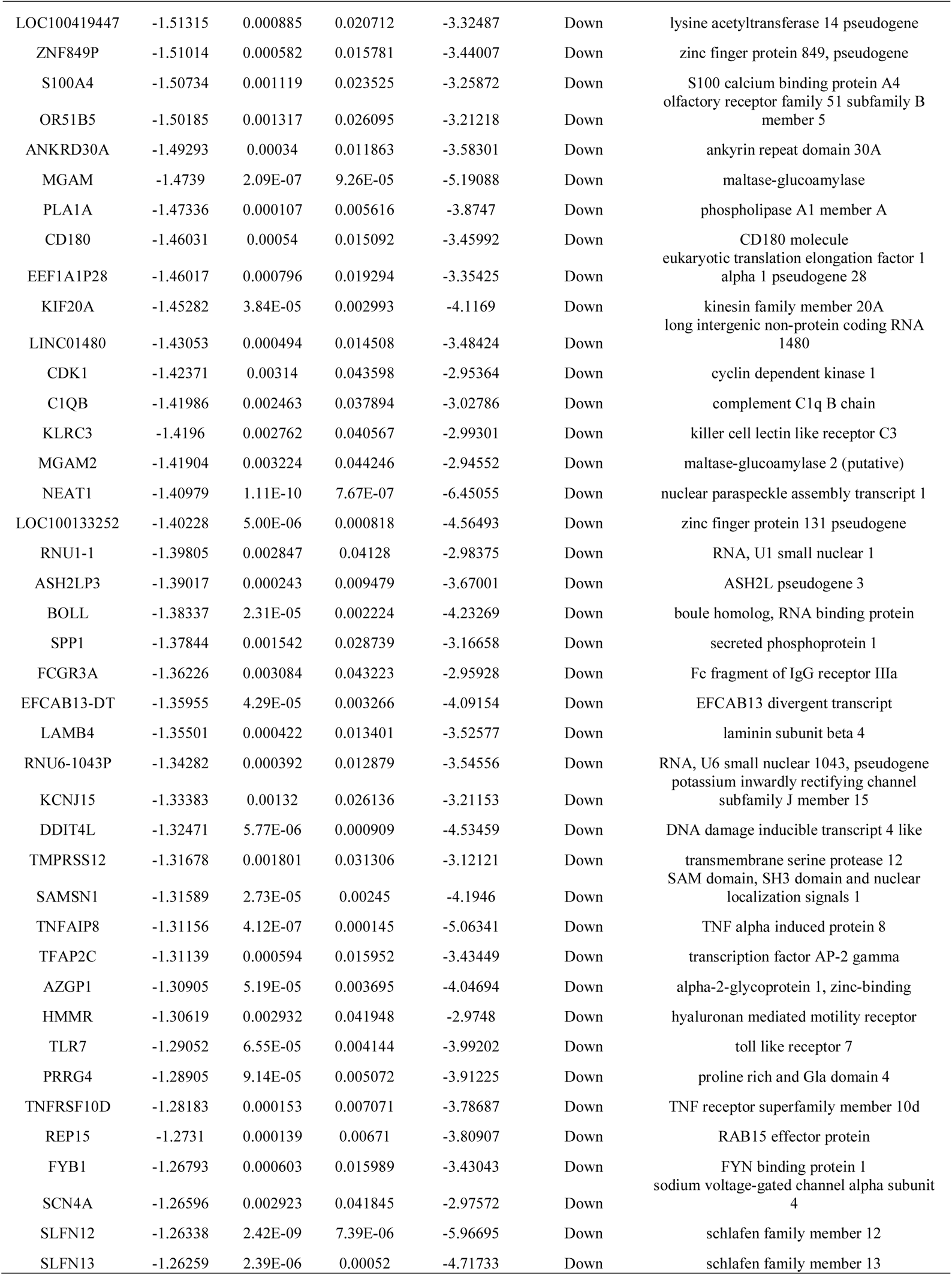
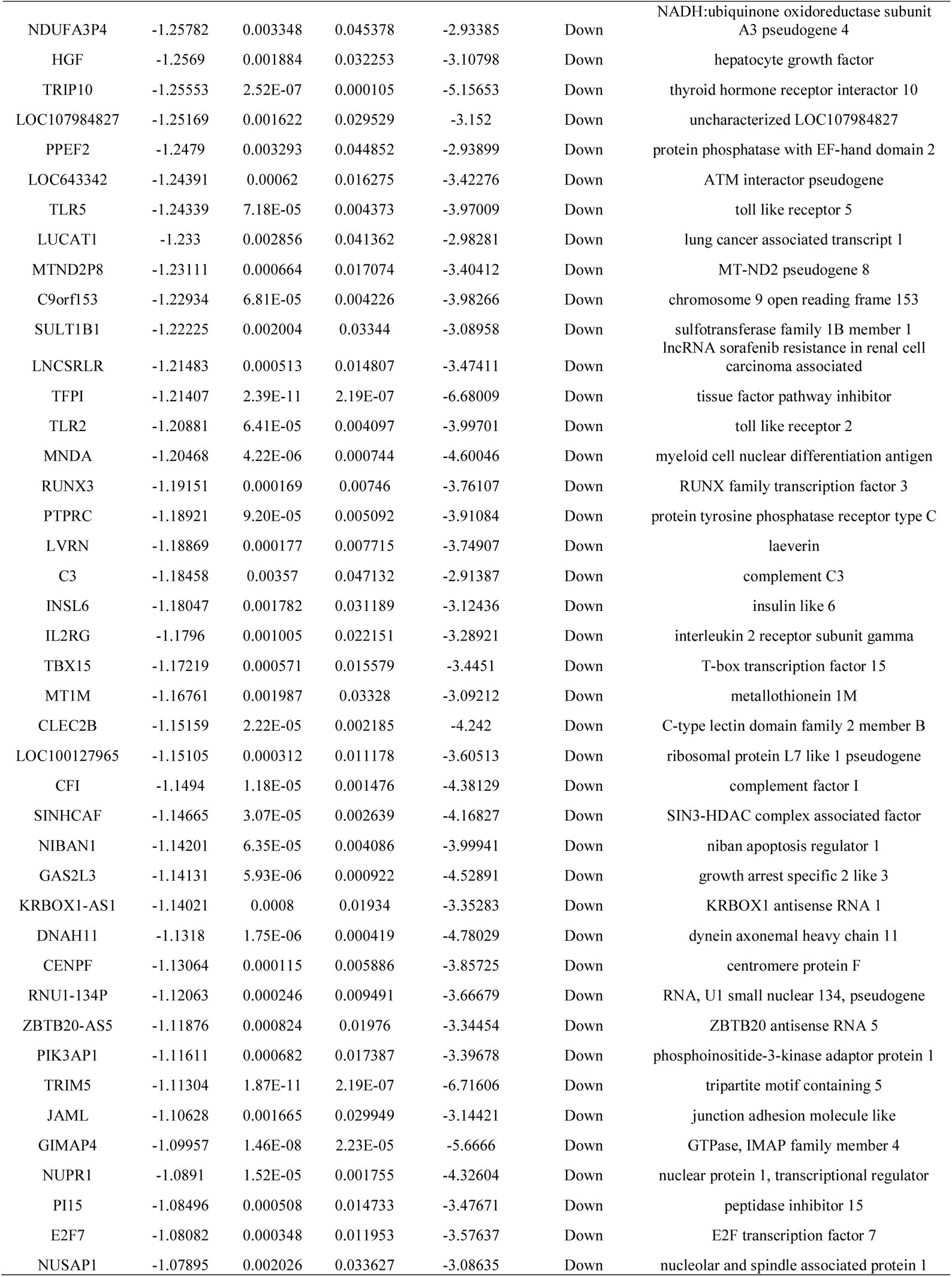
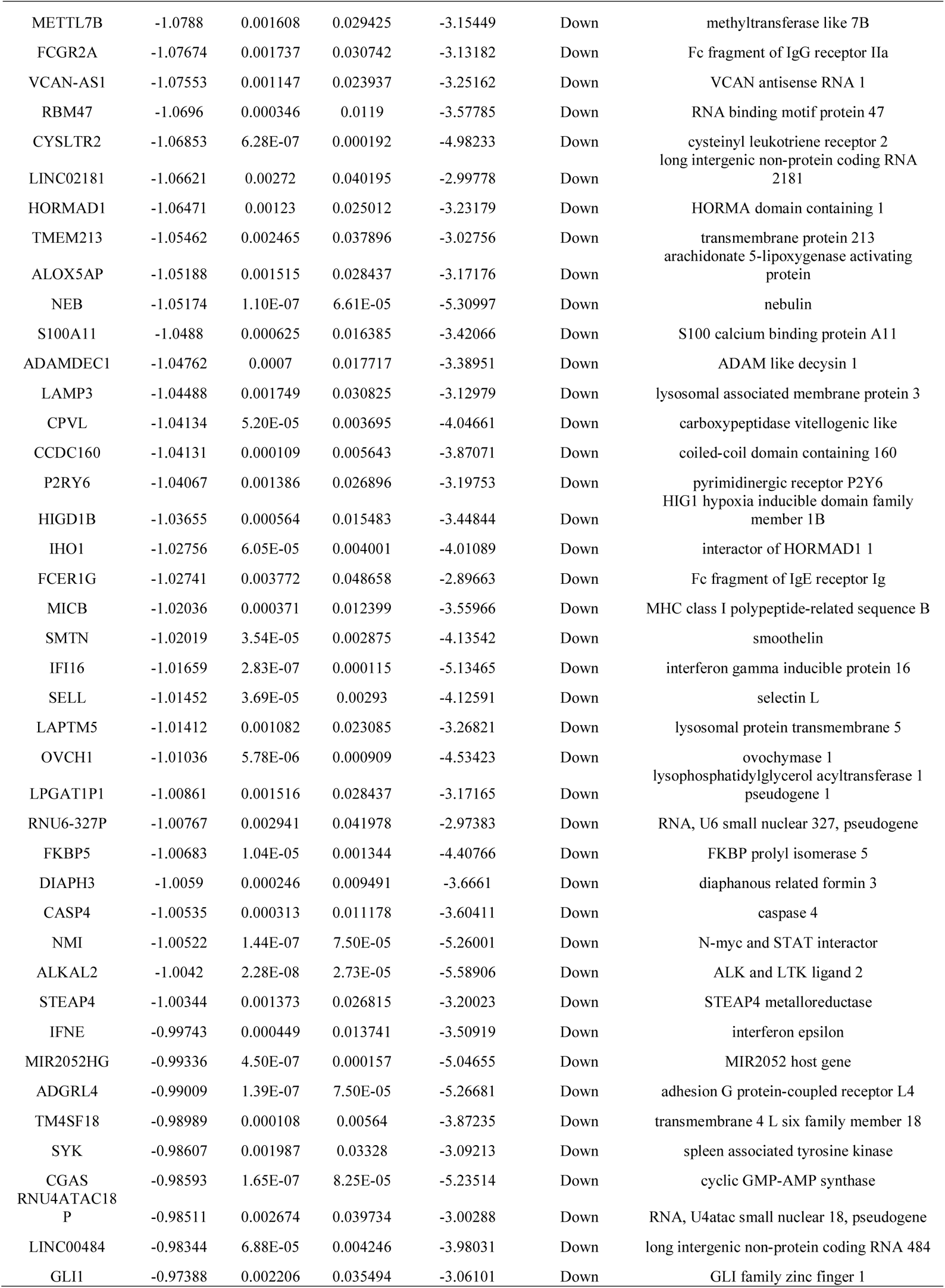
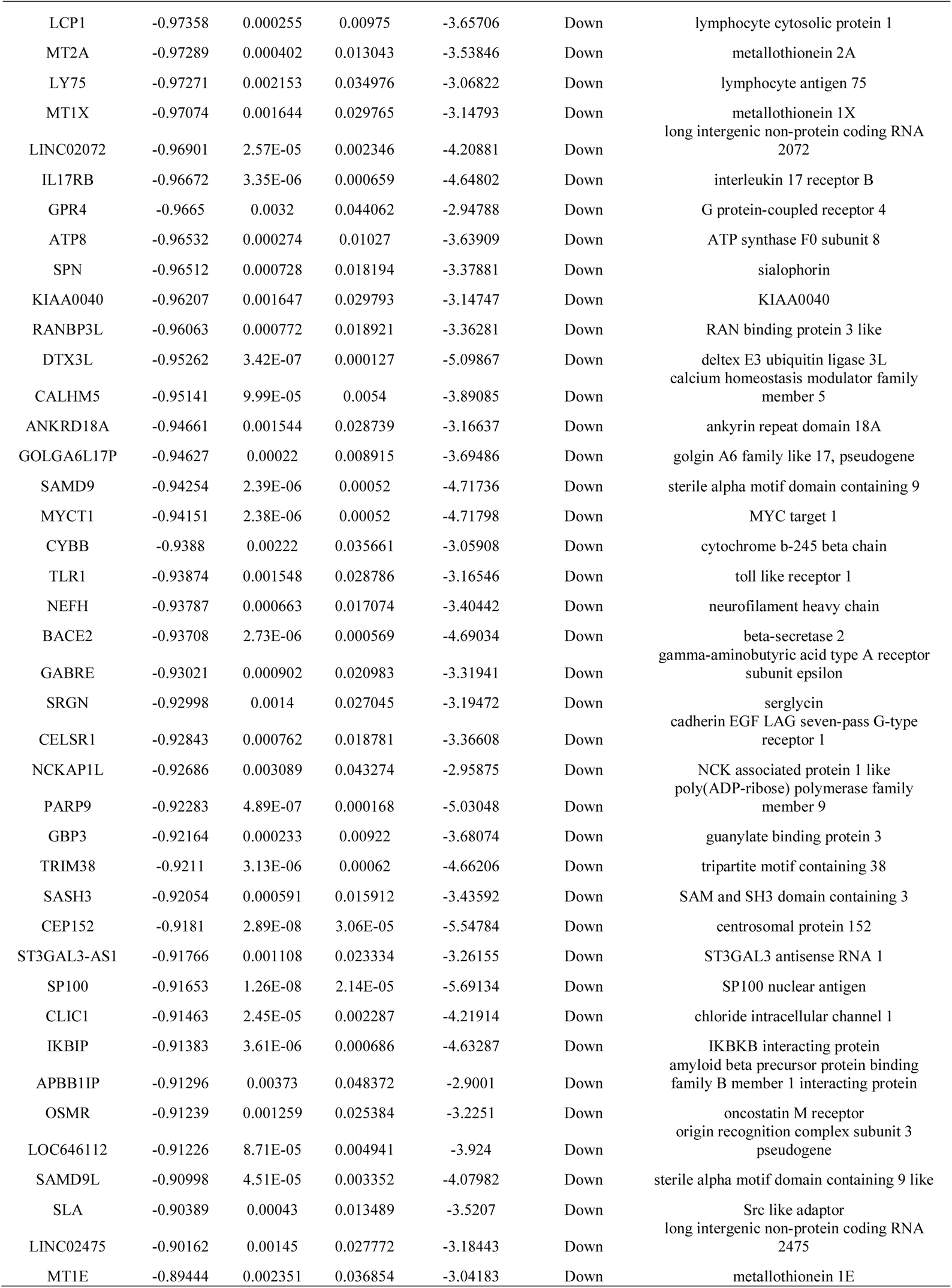
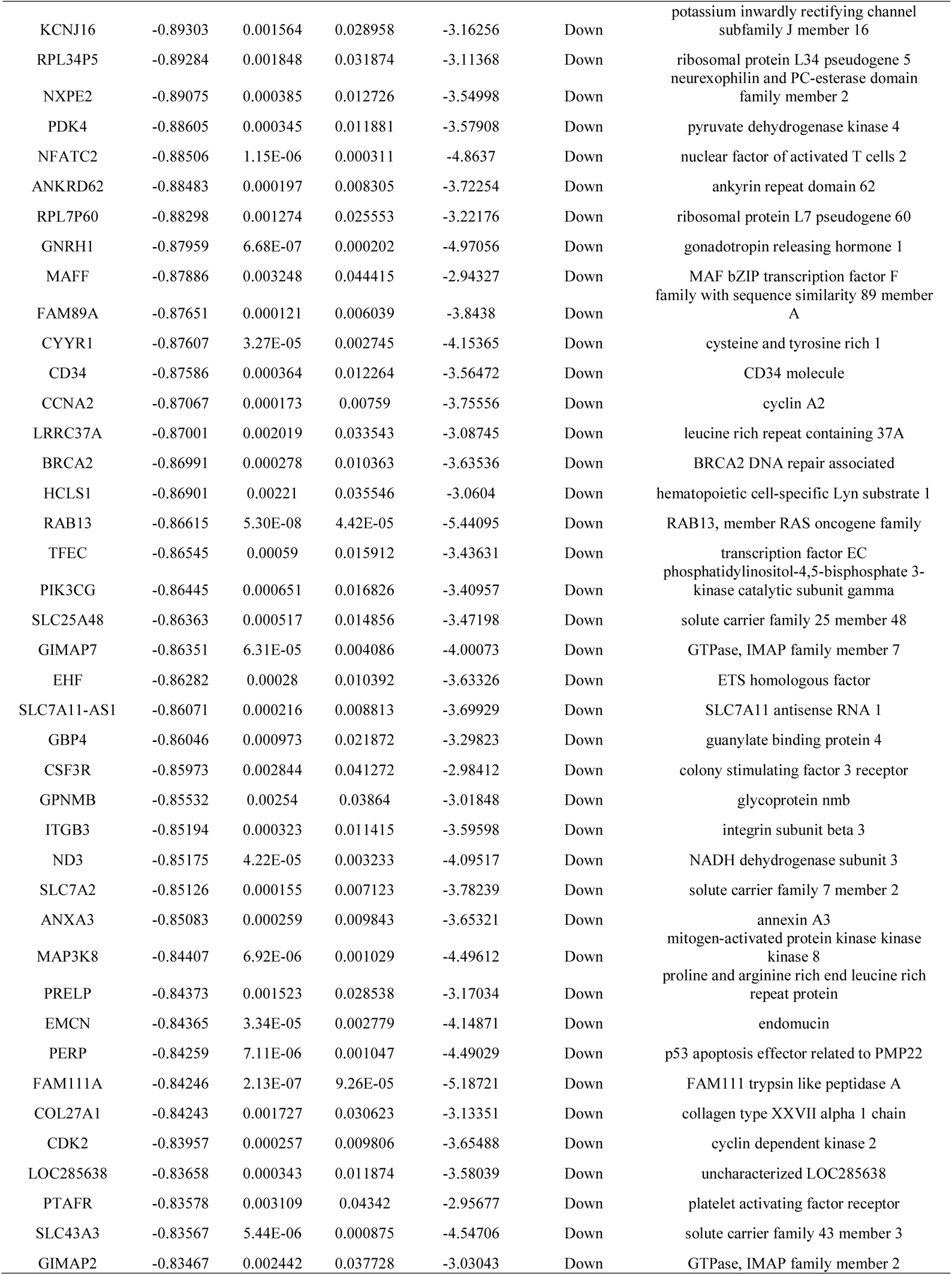
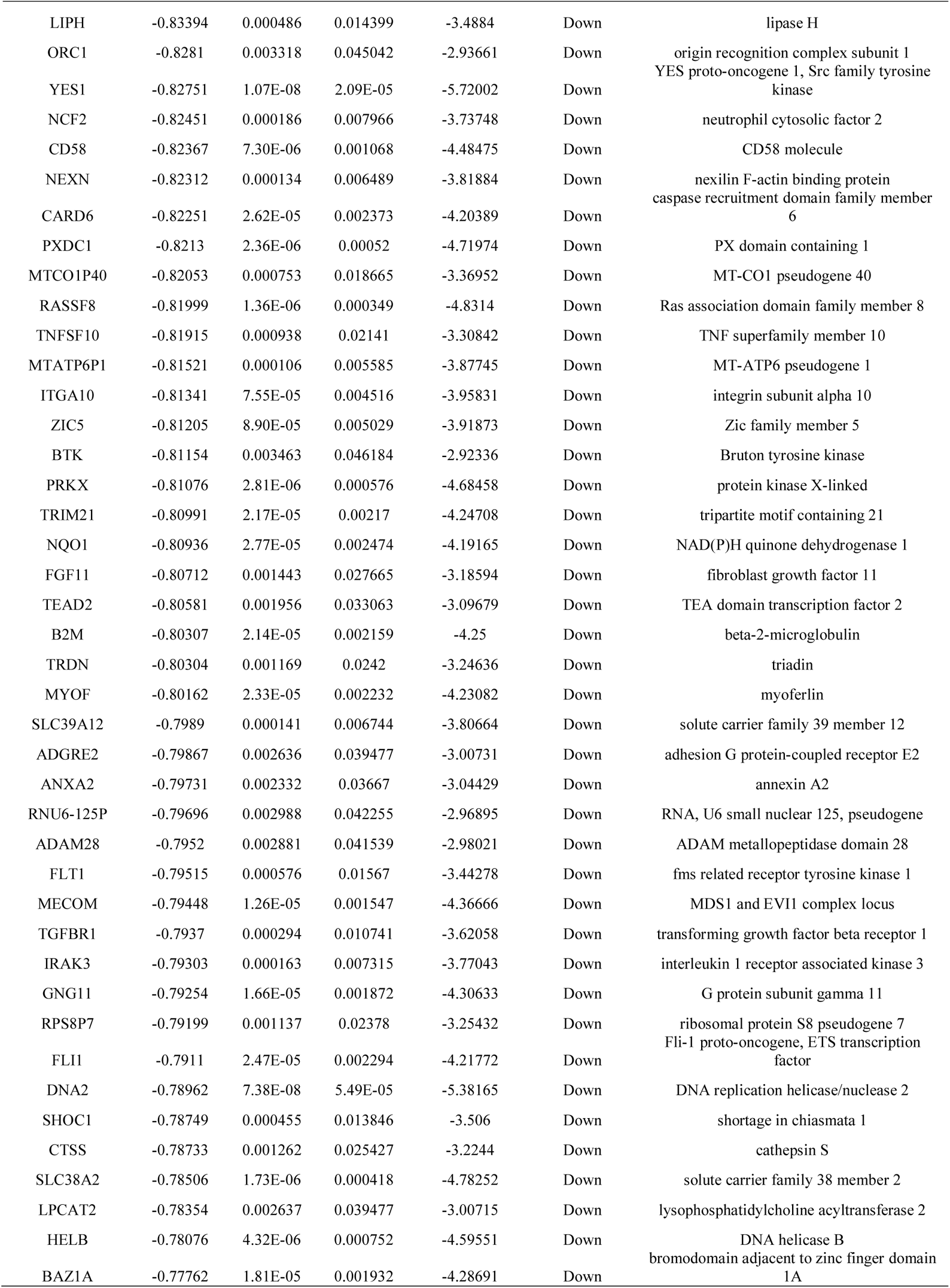
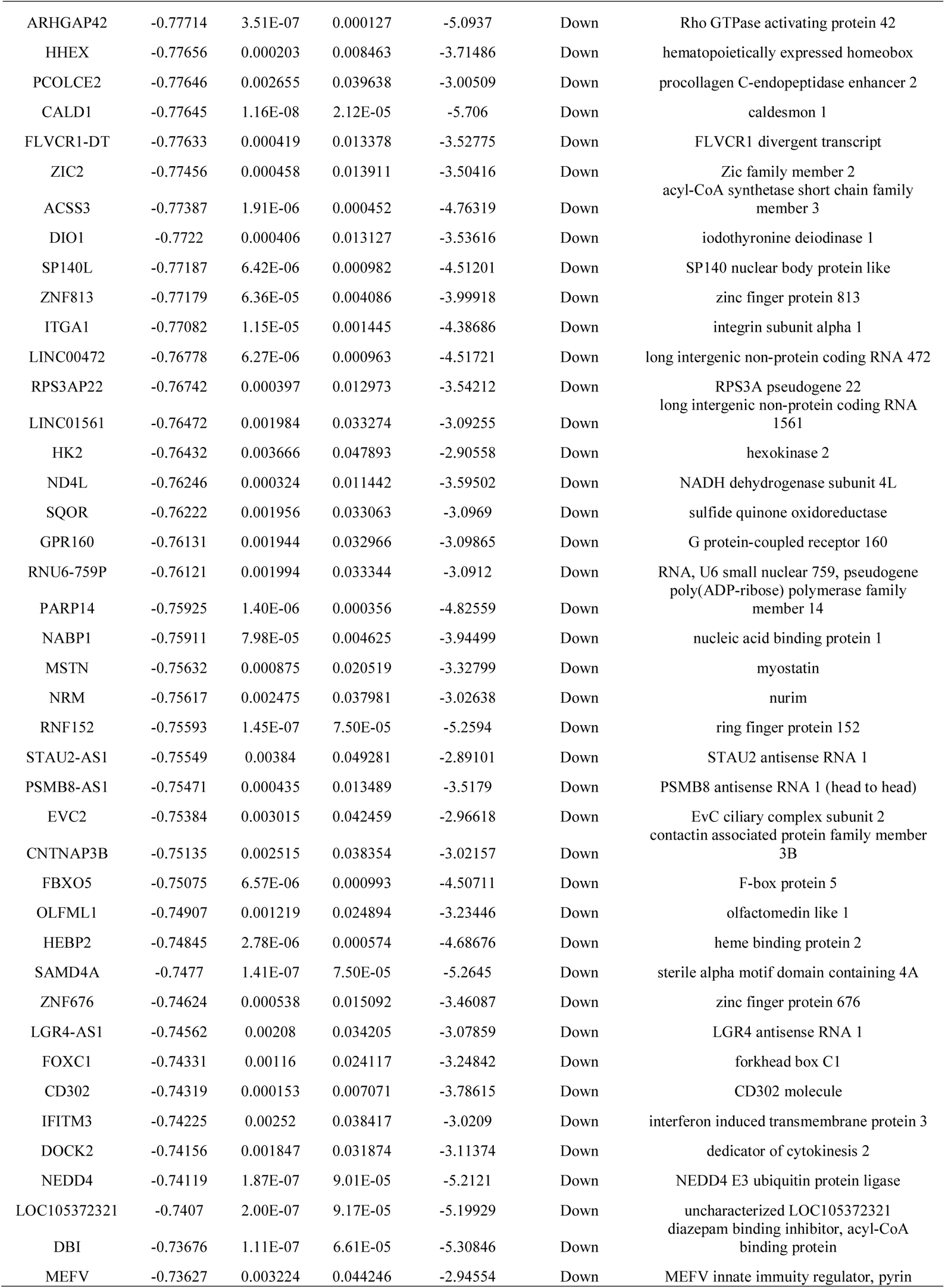
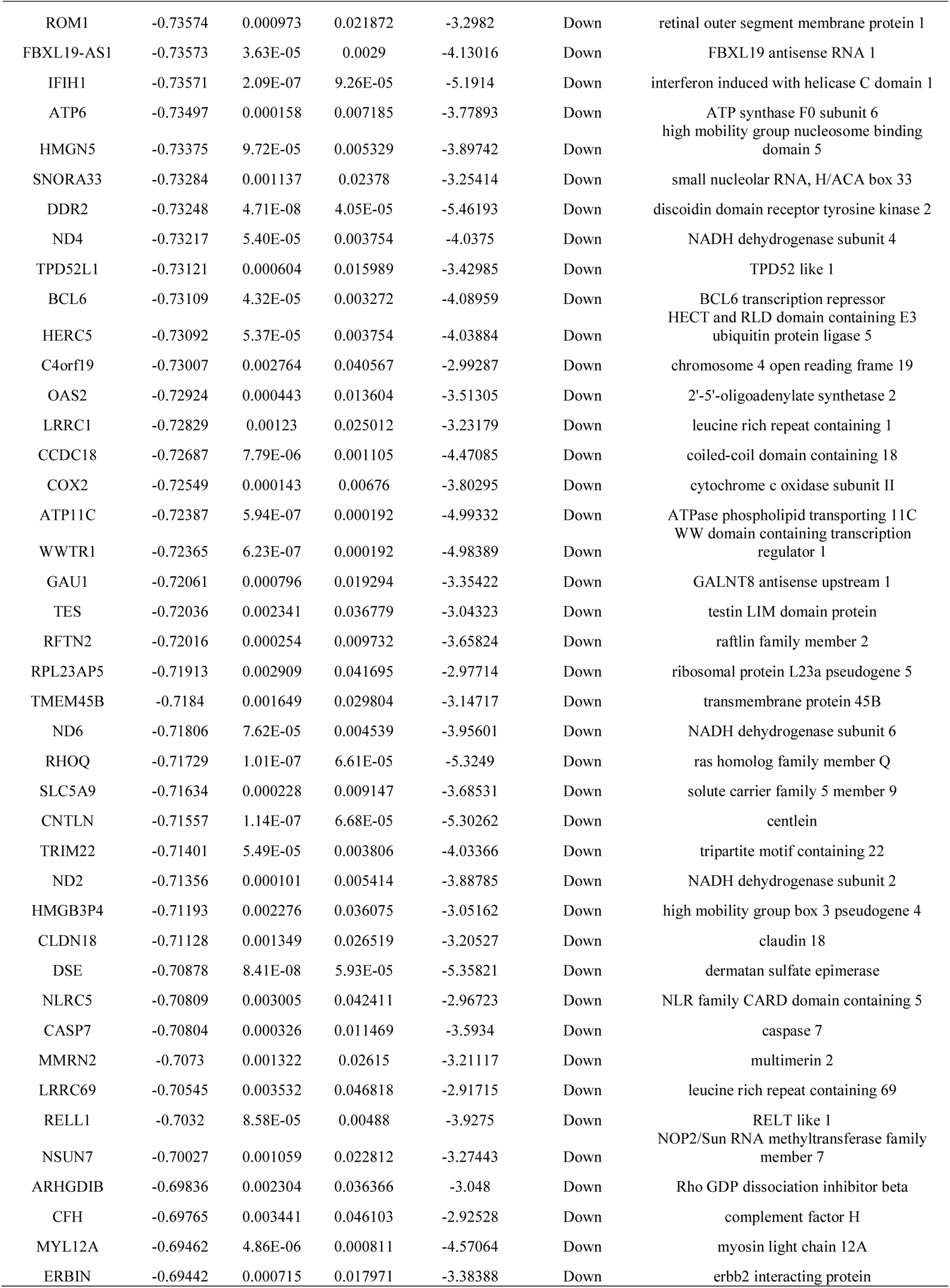
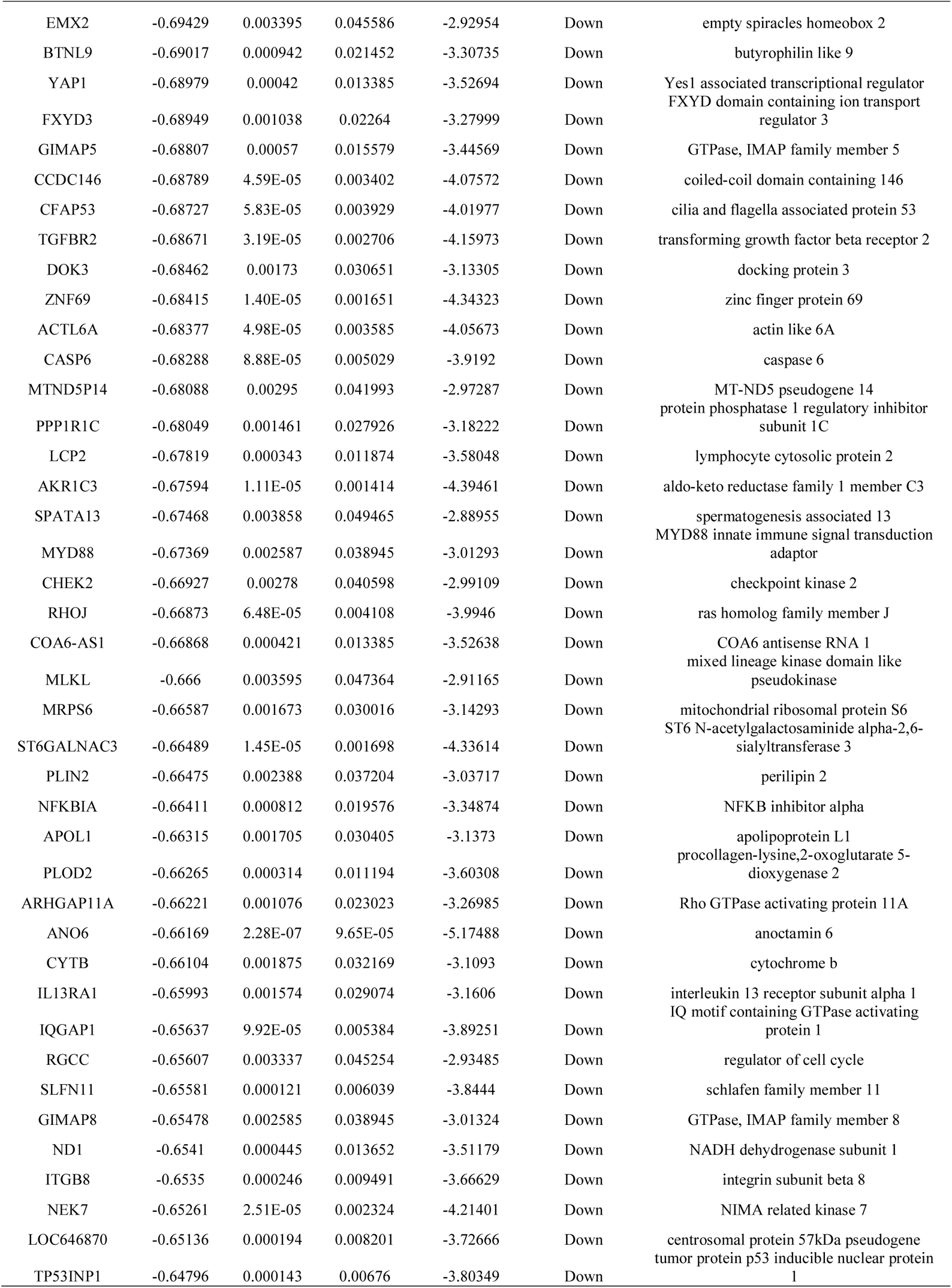
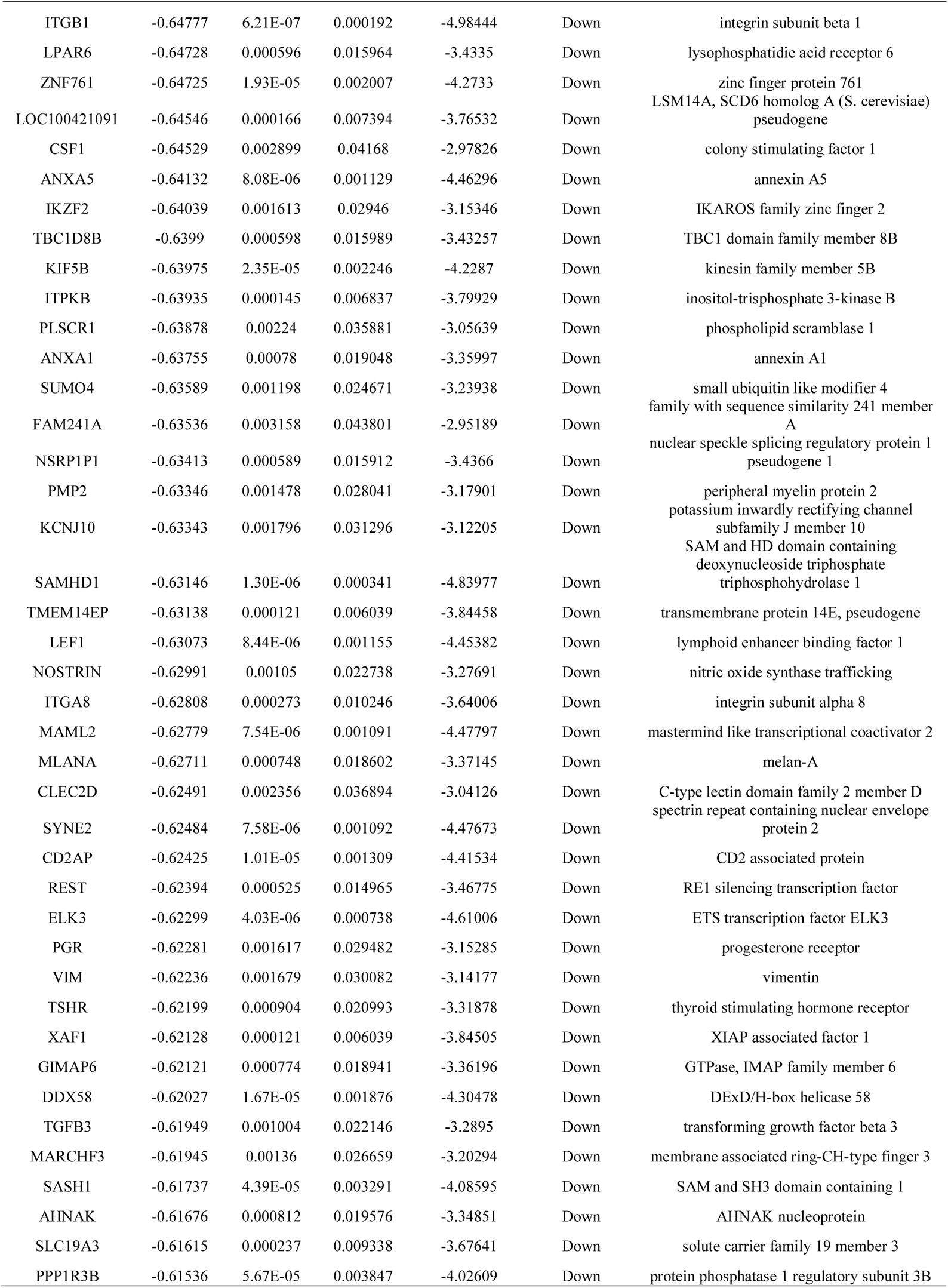
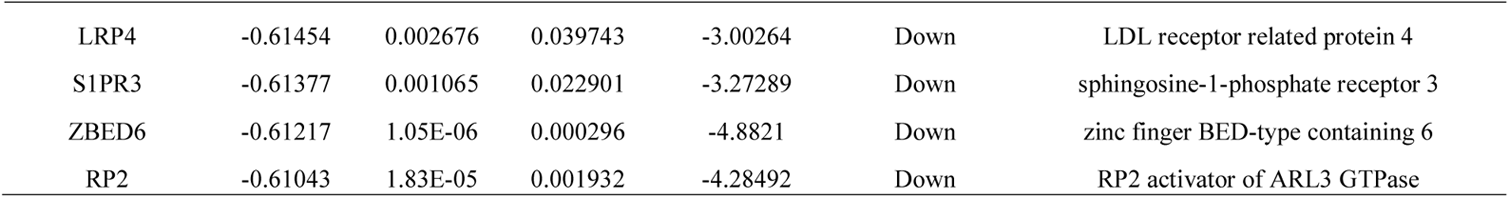
The statistical metrics for key differentially expressed genes (DEGs)

### GO and pathway enrichment analyses of DEGs

In order to better understand the biological function of DEGs, we conducted GO and REACTOME pathway enrichment analysis by g:Profiler. GO results showed that up and down regulated genes significantly enriched in nervous system development, cell communication and response to stimulus of BP, cell junction, membrane, cell periphery and cytoplasm of CC, and transporter activity, protein binding, identical protein binding and molecular transducer activity of MF (Table 2). Moreover, REACTOME pathway enrichment analysis showed that the up and down regulated genes were enriched in neuronal system, transmission across chemical synapses, immune system and cytokine signaling in immune system (Table 3).

**Table 2.**
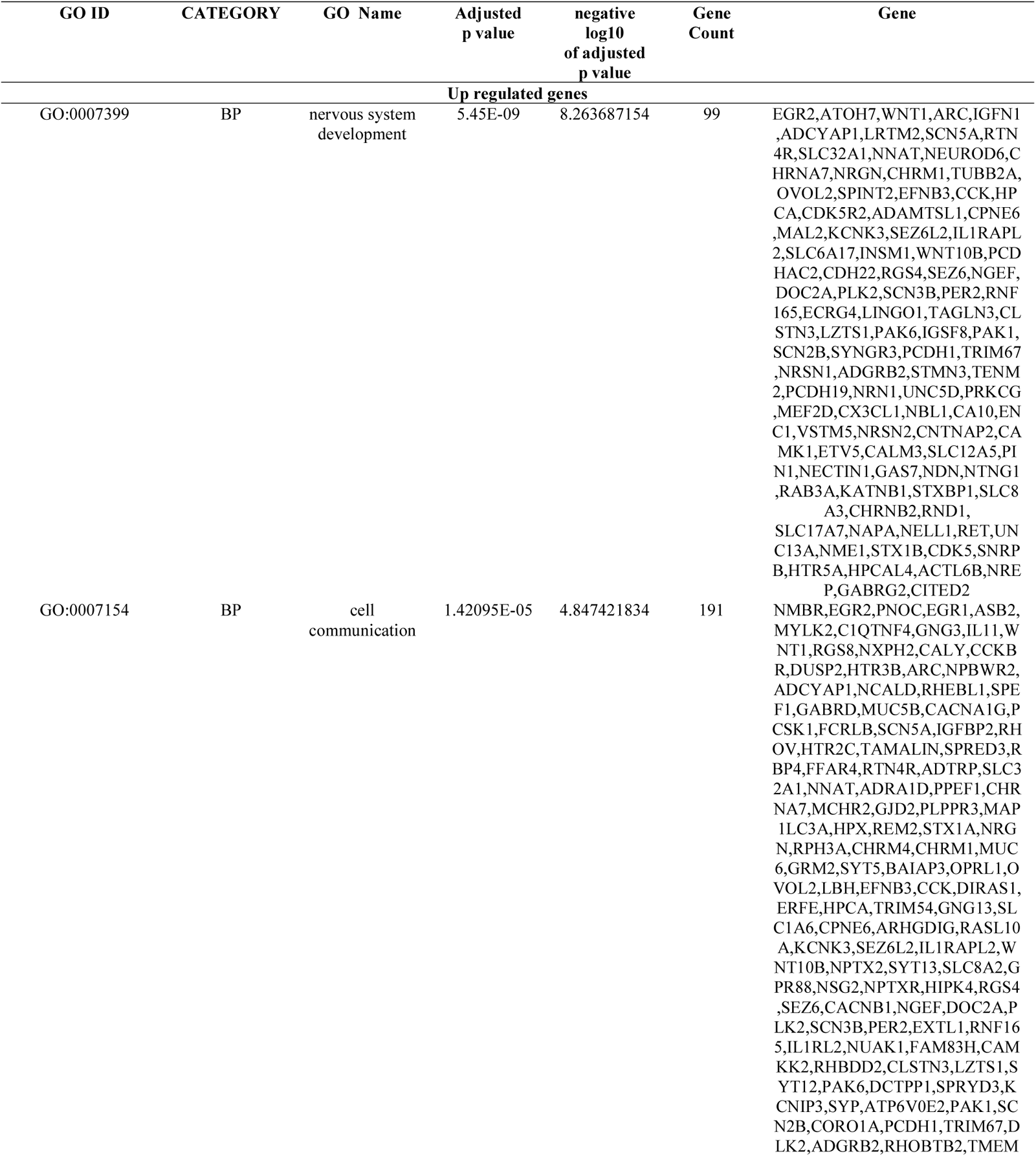
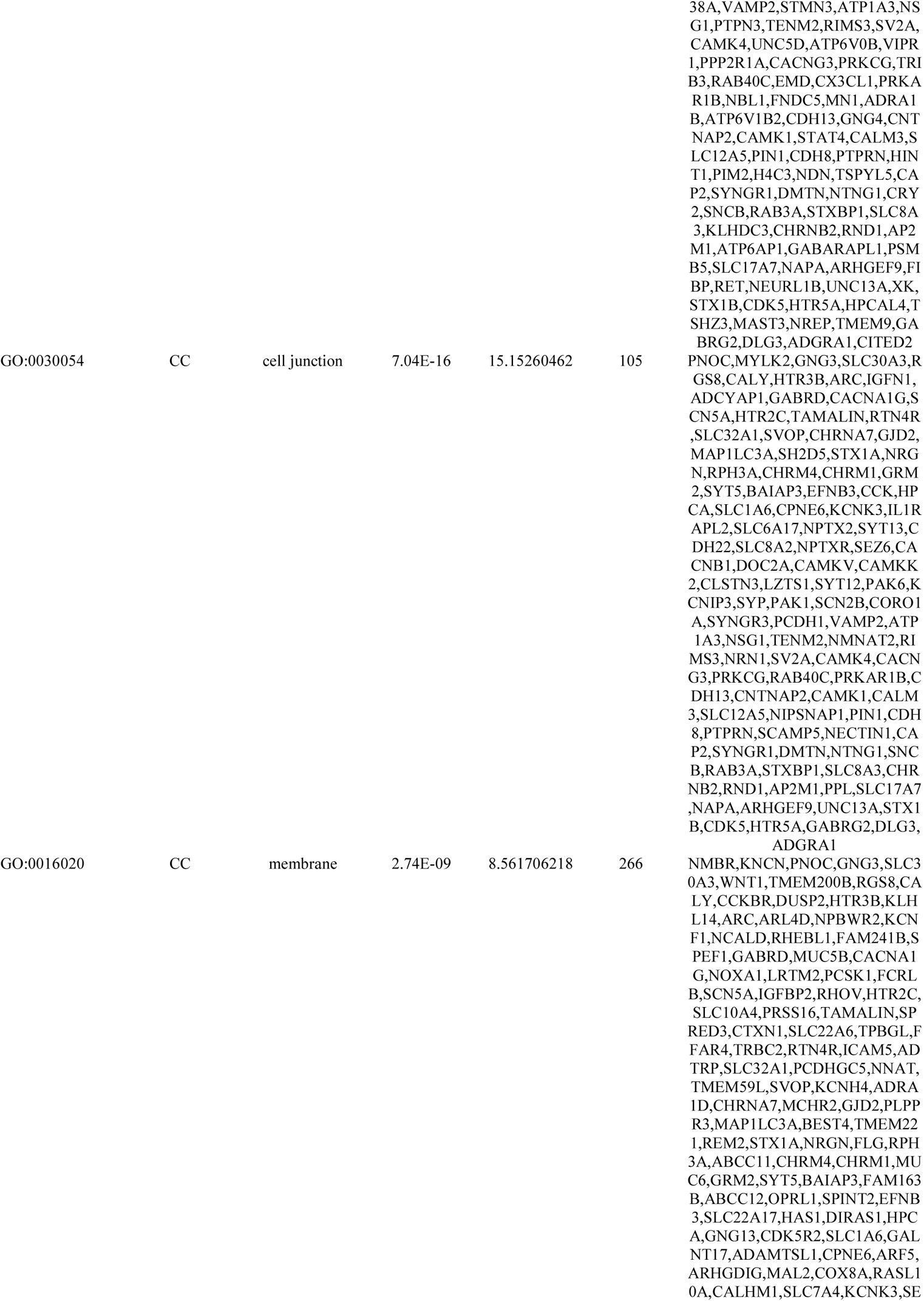
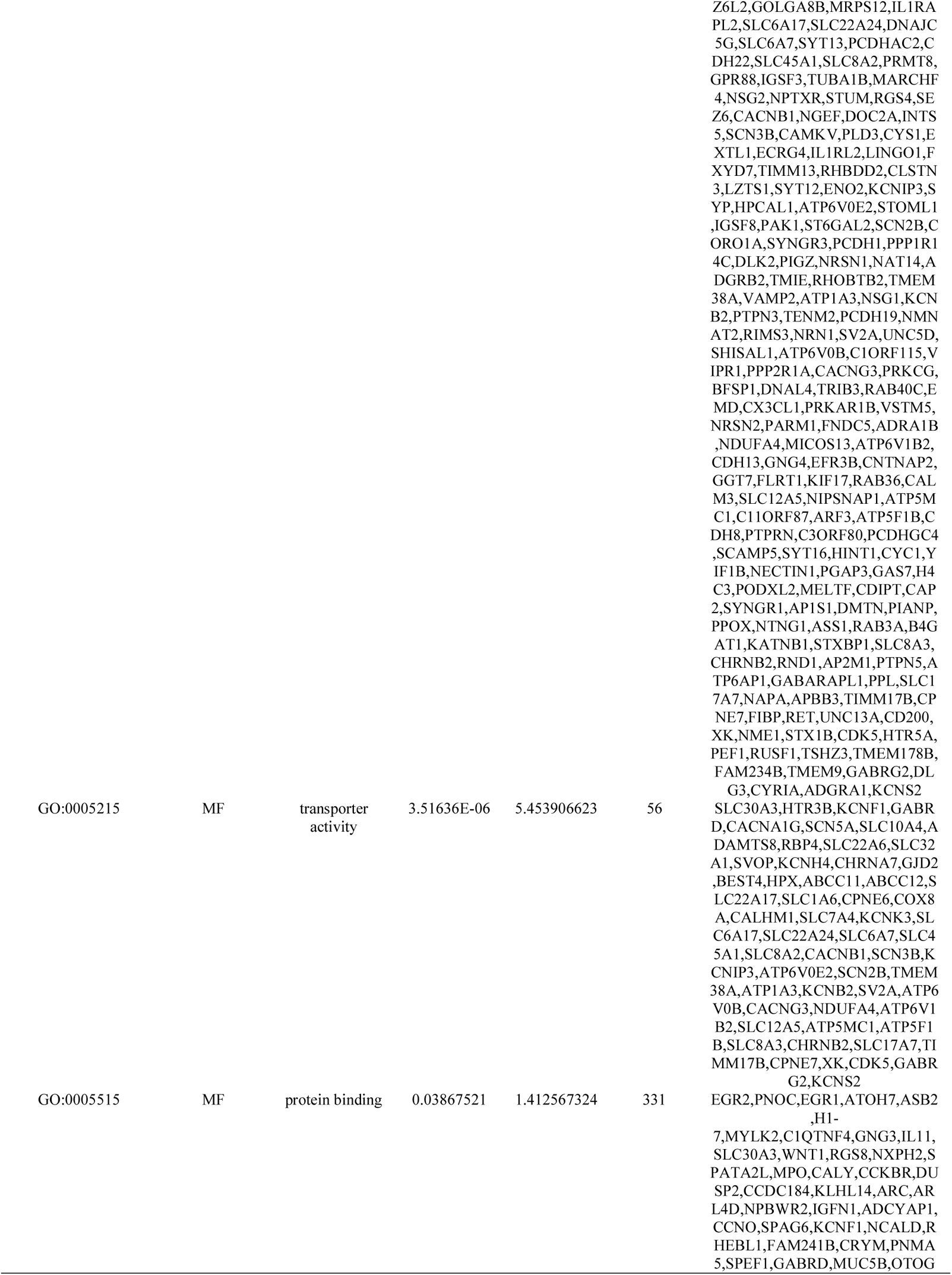
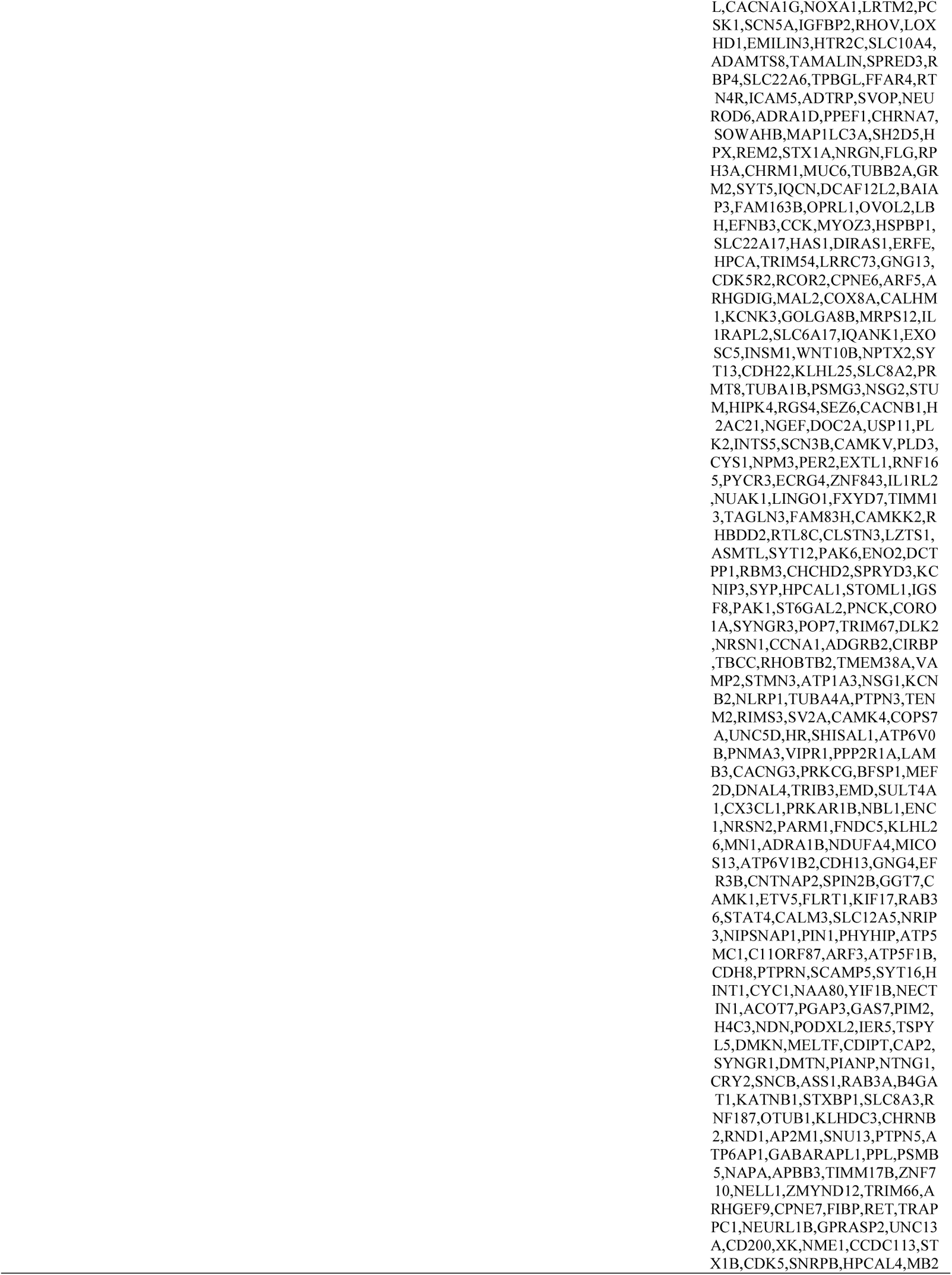
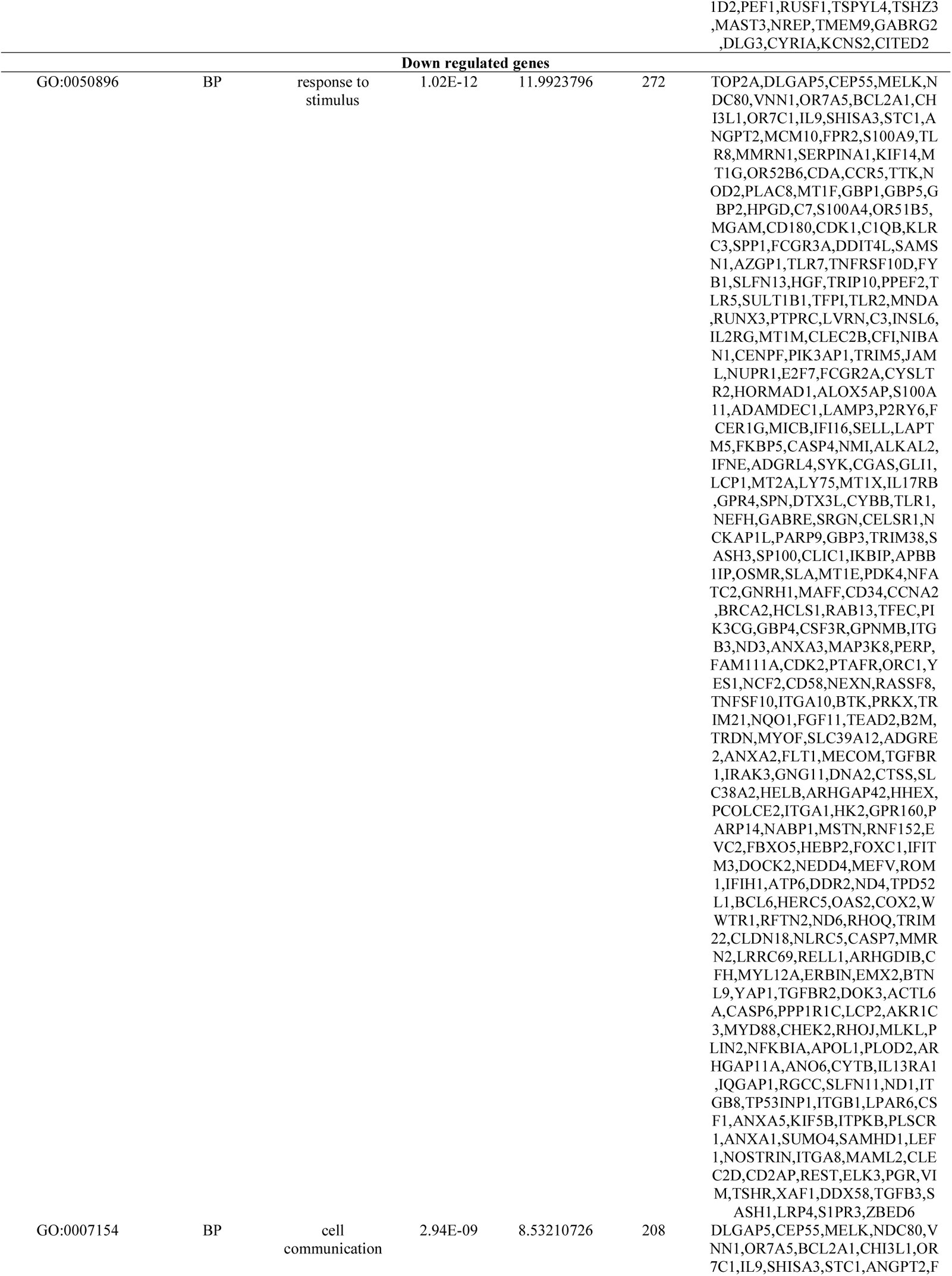
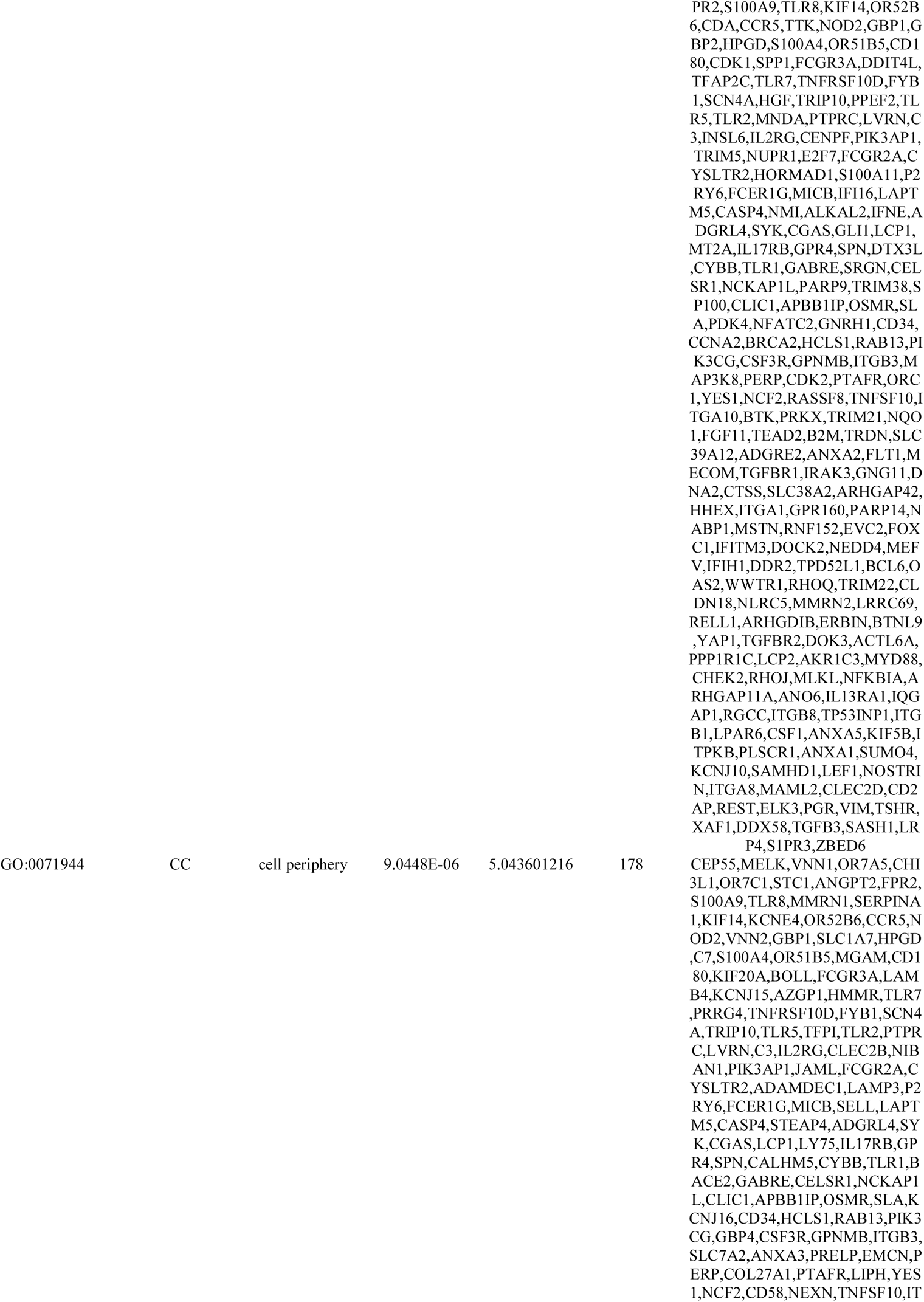
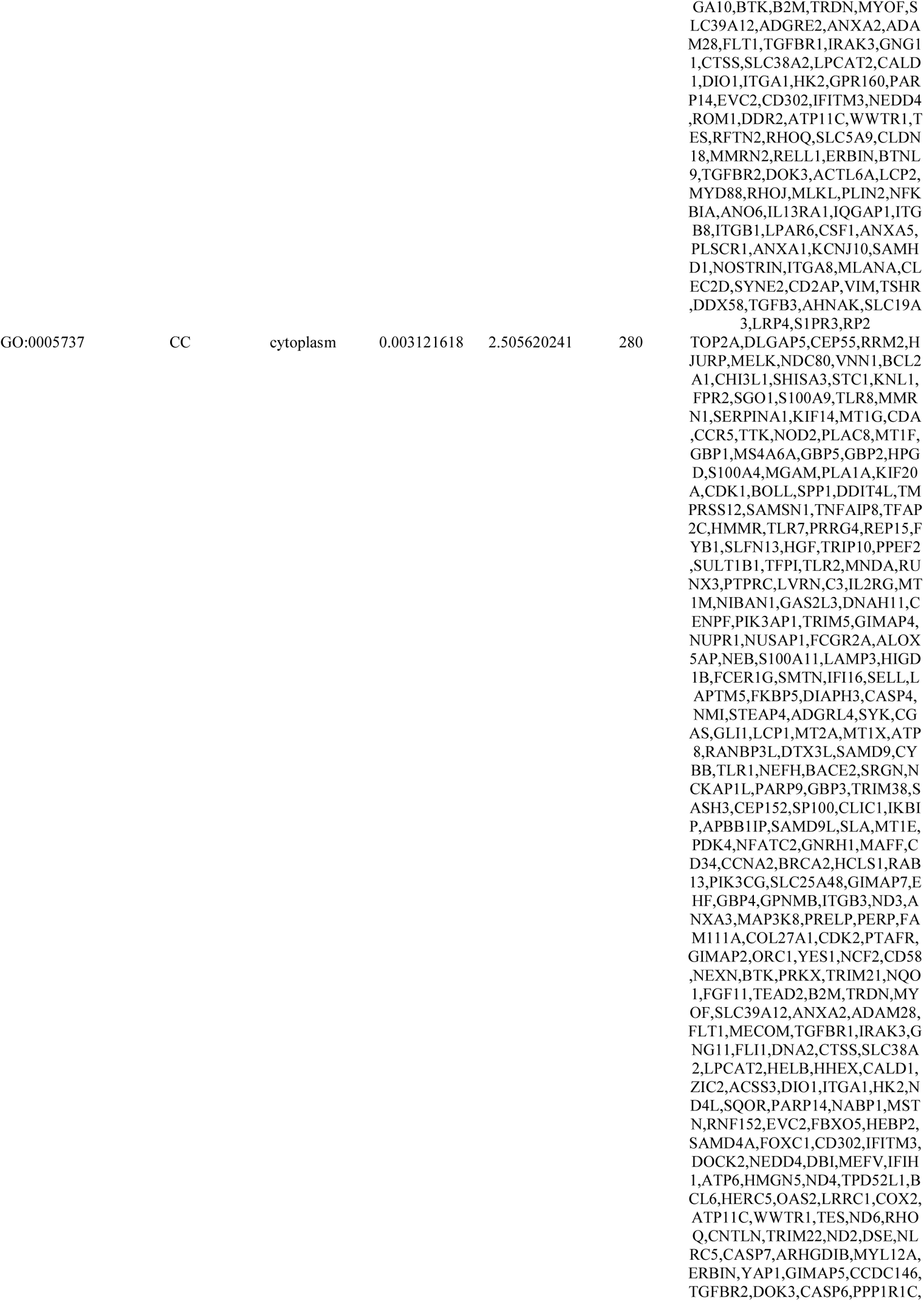
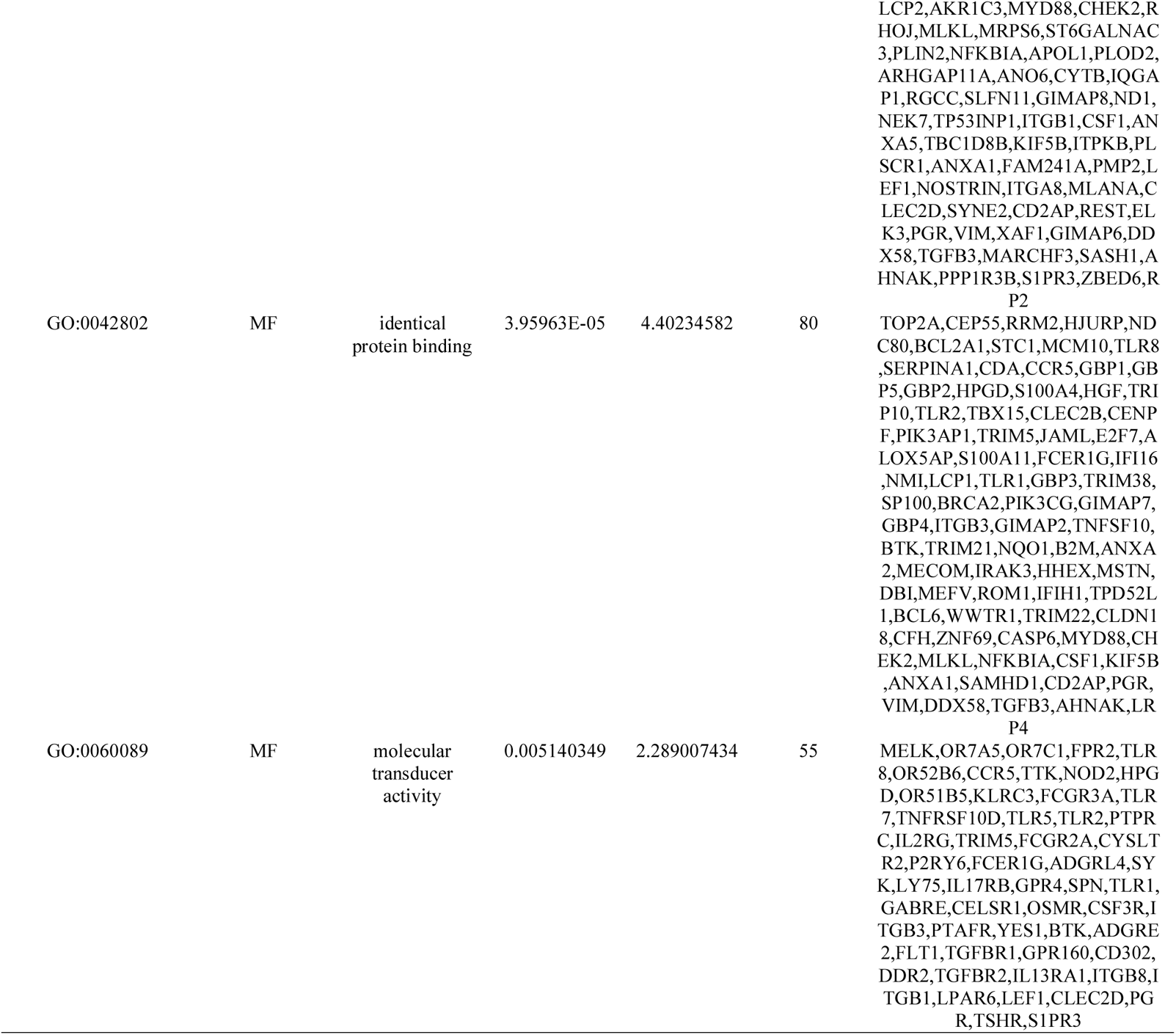
The enriched GO terms of the up and down regulated differentially expressed genes

**Table 3.**
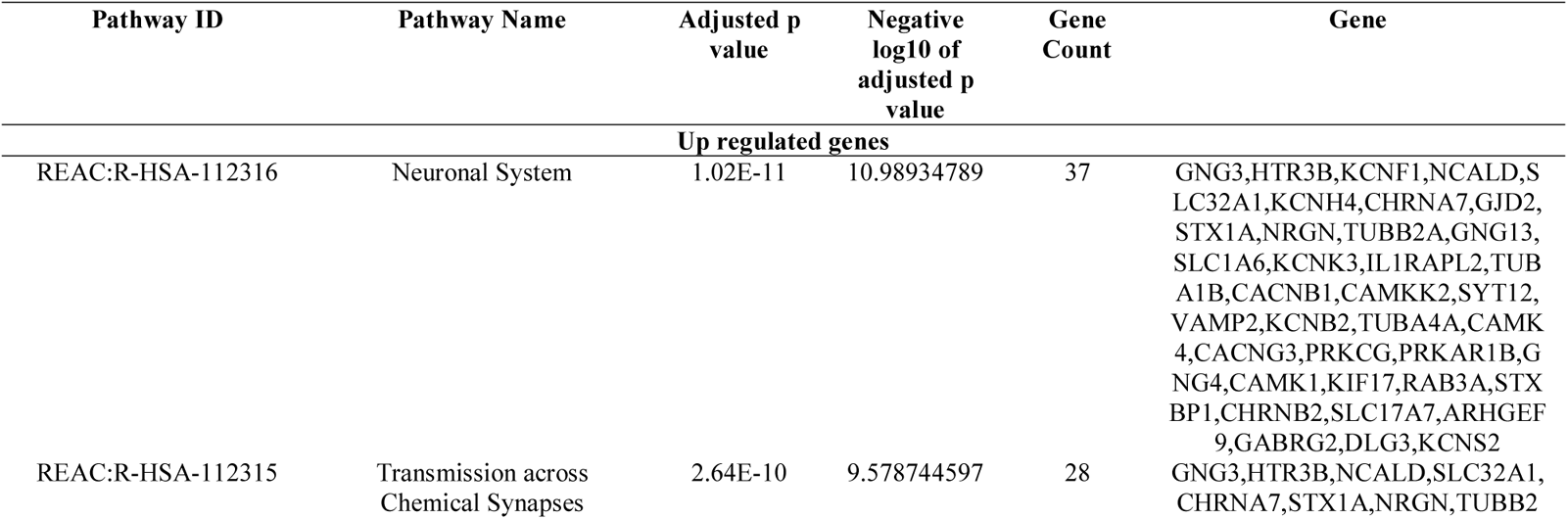
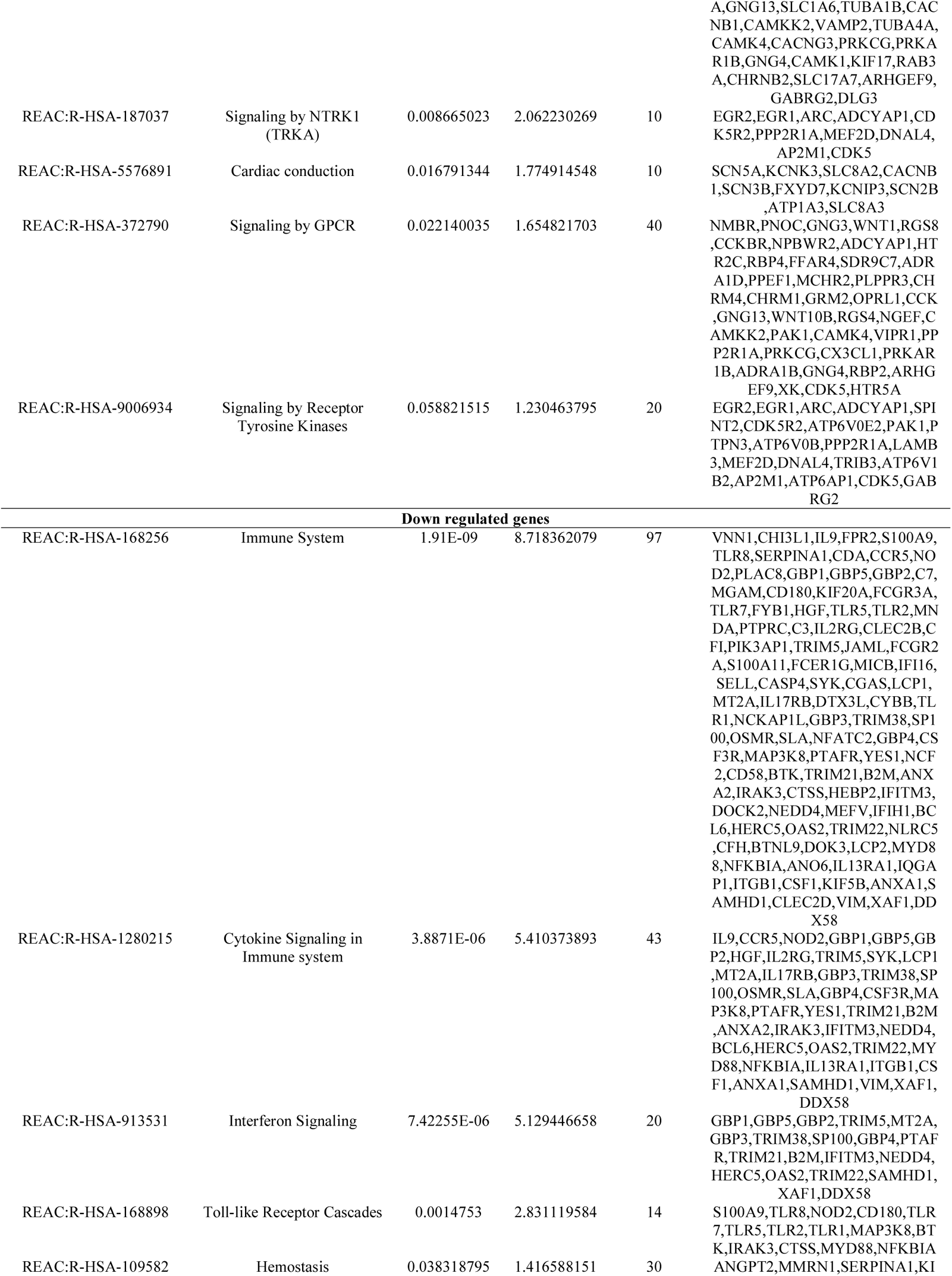
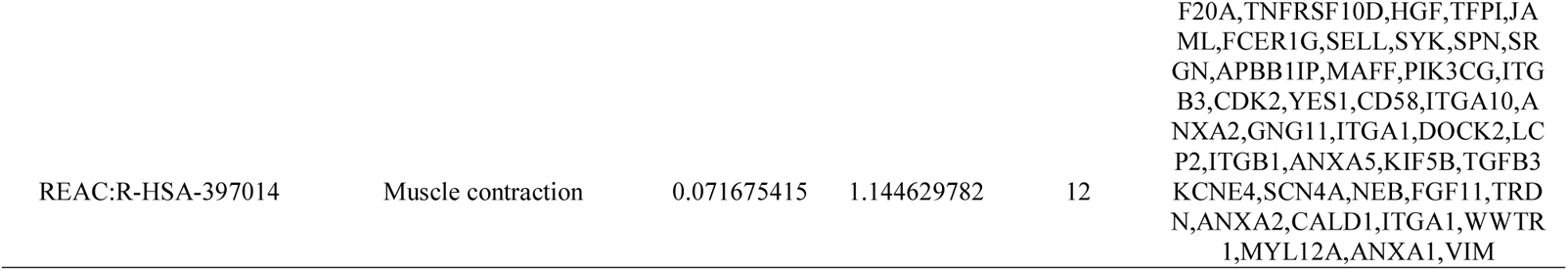
The enriched pathway terms of the up and down regulated differentially expressed genes

### Construction of the PPI network and module analysis

To investigate the molecular mechanism of PD from a systematic perspective, PPI network was built to examine the relationship between proteins. PPI network was built by HIPPIE interactome for DEGs. There were 4092 nodes and 7138 edges in the visualization network using the Cyctoscape (Fig. 3). Based on the high node degree, betweenness centrality, stress centrality and closeness centrality the top hub genes, including OTUB1, PPP2R1A, AP2M1, PIN1, USP11, CDK2, IQGAP1, NEDD4, VIM and CDK1, were identified in the PPI network (Table 4). PEWCC1was used to identify the significant cluster modules in the PPI network and the top 2 modules were selected (Fig. 4A and 4B). Following GO and REACTOME pathway screening, the module 1 (7 nodes and 15 edges) was revealed to be associated with neuronal system and nervous system development and the module 2 (20 nodes and 41 edges) was revealed to be associated with immune system, muscle contraction, signaling by NTRK1 (TRKA), response to stimulus and cell communication.

**Fig. 3.**
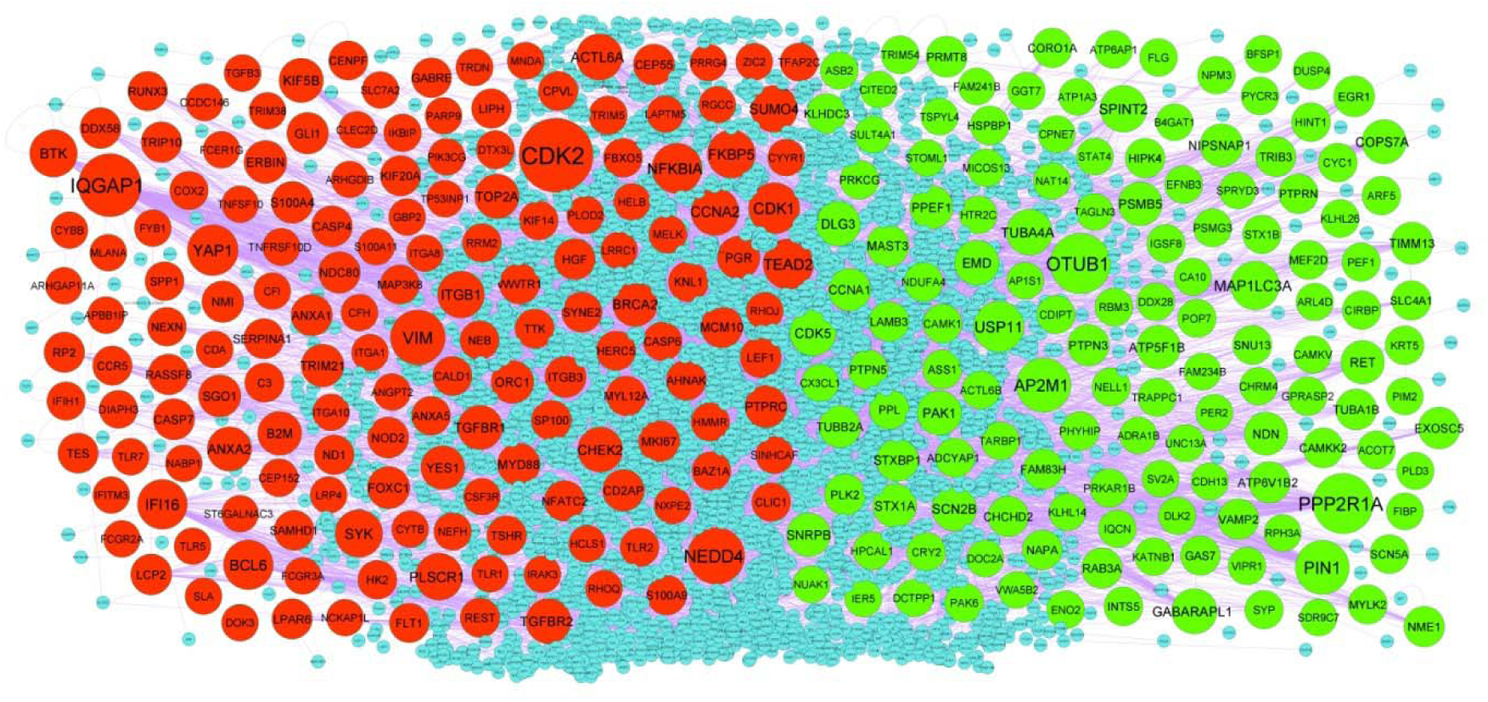
PPI network of DEGs. Up regulated genes are marked in green; down regulated genes are marked in red

**Fig. 4.**
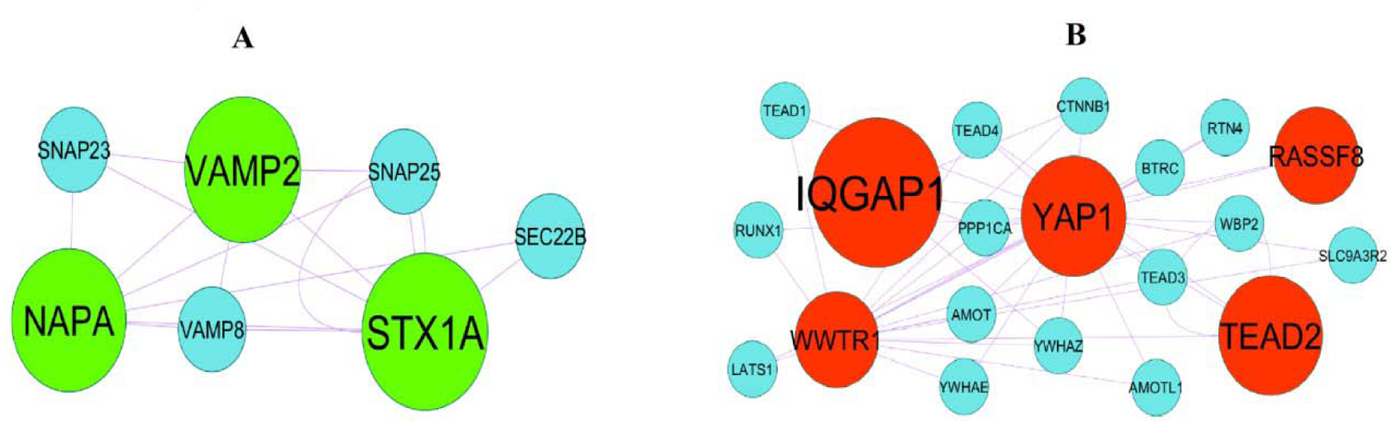
Modules of isolated form PPI of DEGs. (A) The most significant module was obtained from PPI network with 7 nodes and 15 edges for up regulated genes (B) The most significant module was obtained from PPI network with 20 nodes and 41 edges for down regulated genes. Up regulated genes are marked in green; down regulated genes are marked in red

**Table 4.**
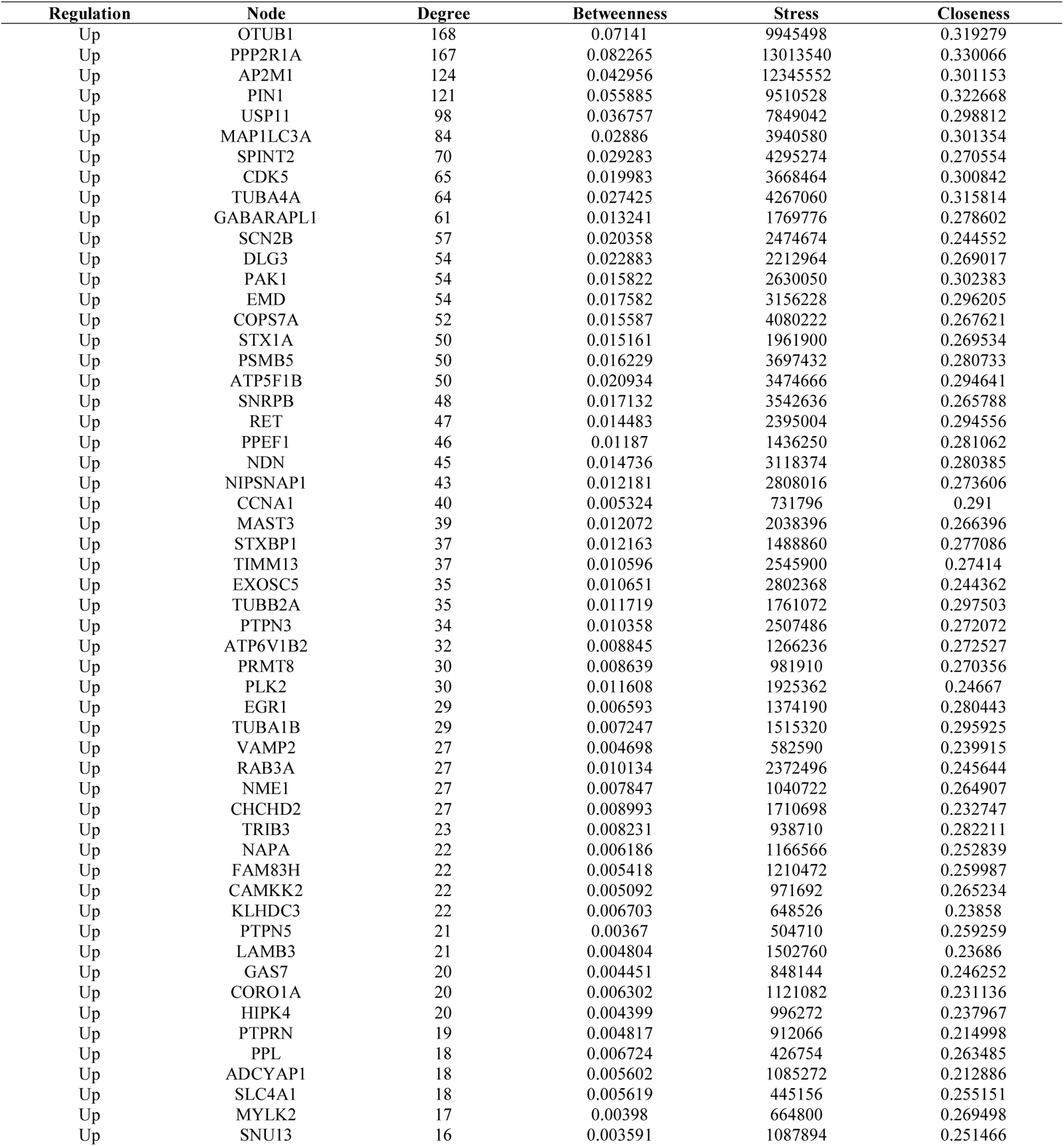
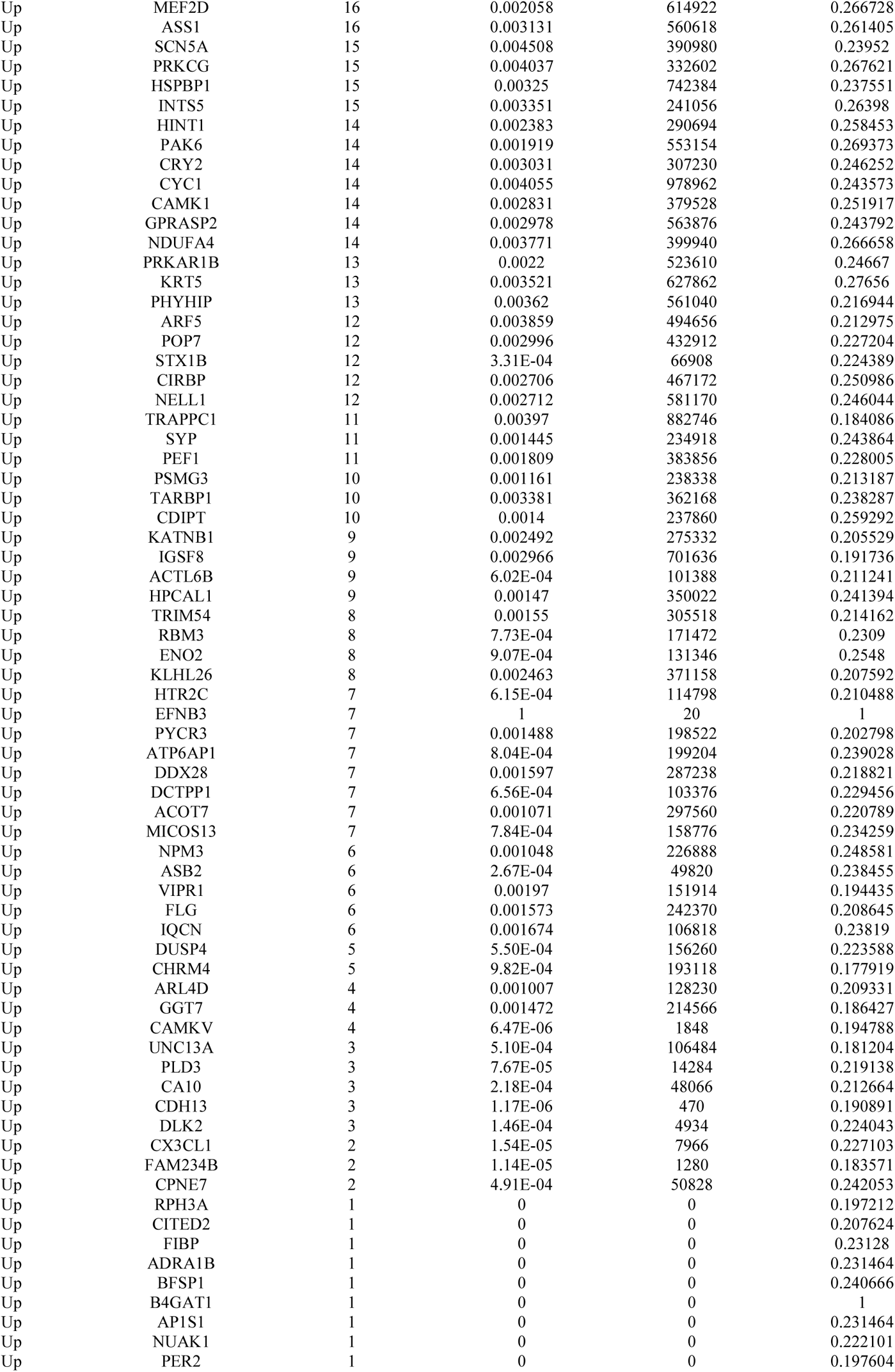
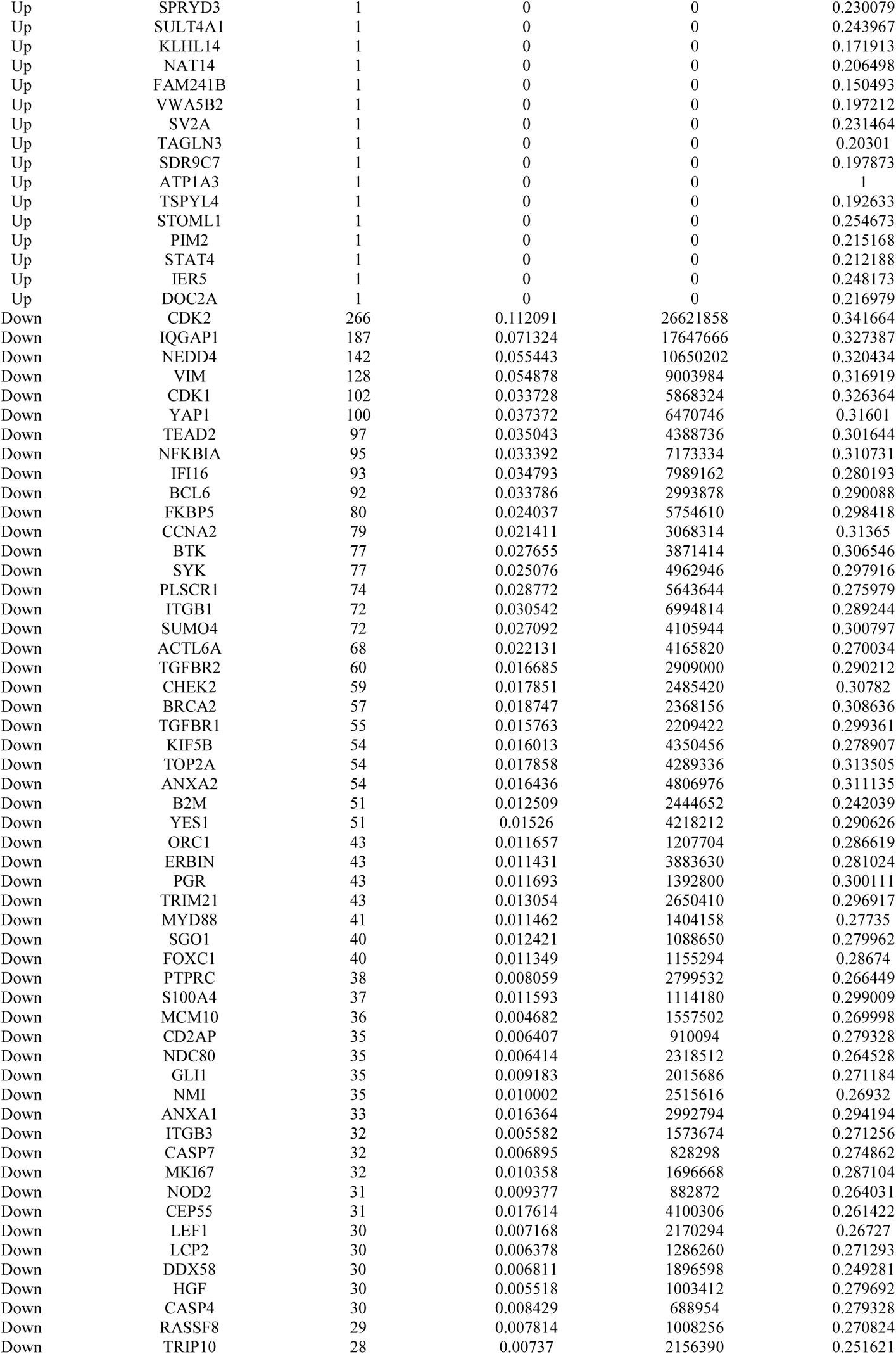
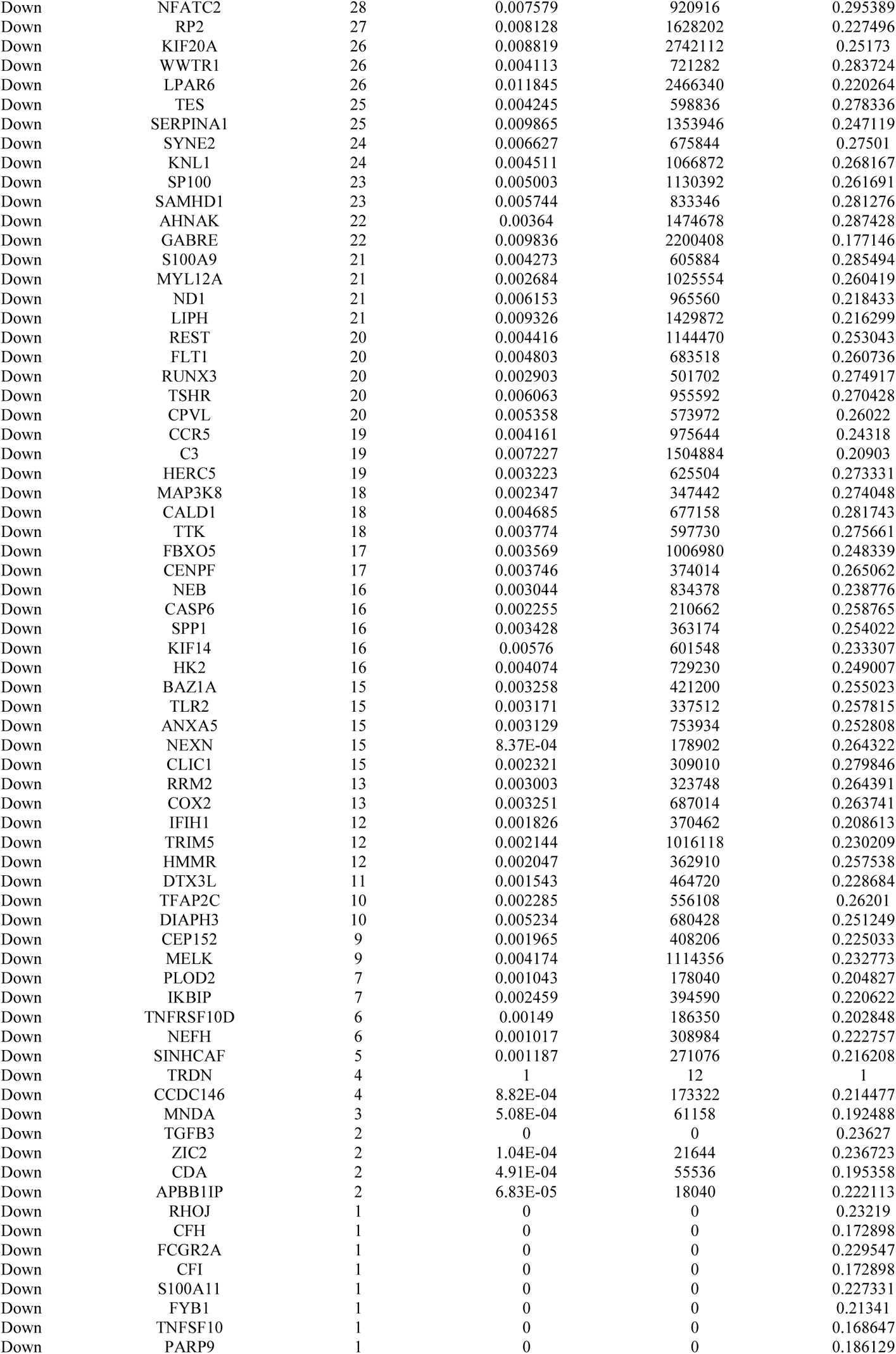
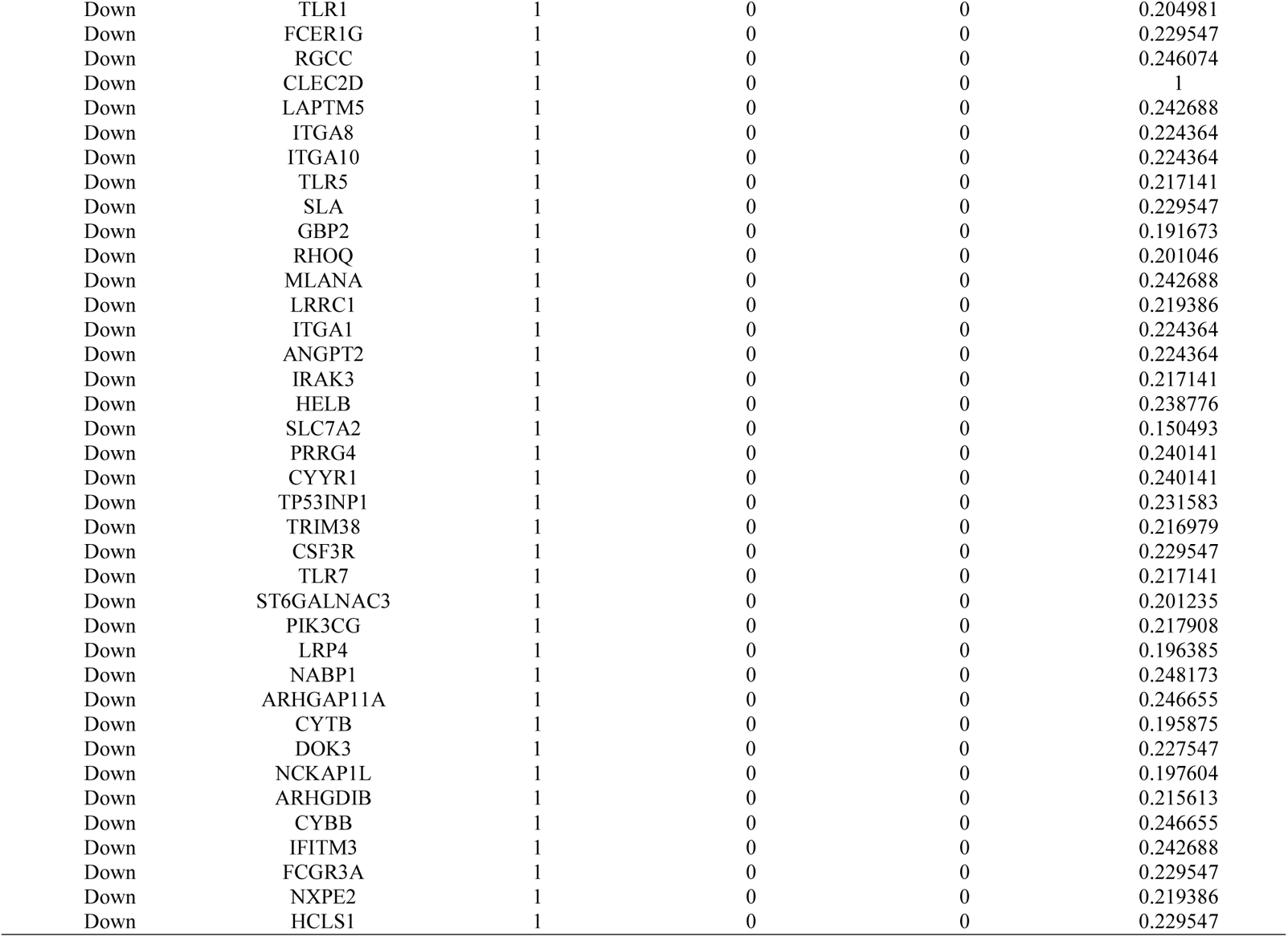
Topology table for up and down regulated genes.

### miRNA-hub gene regulatory network construction

miRNA-hub gene regulatory network was built by miRNet for hub genes. The miRNA-hub gene regulatory network of hub genes was constructed with 2432 (miRNA: 2135; hub gene: 297) nodes and 14589 edges (Fig. 5). AP2M1 was targeted by 69 miRNAs (ex; hsa-mir-3911), PIN1 was targeted by 56 miRNAs (ex; hsa-mir-199b-5p), PPP2R1A was targeted by 46 miRNAs (ex; hsa-mir-6779-5p), SCN2B was targeted by 46 miRNAs (ex; hsa-mir-4722-3p), OTUB1 was targeted by 45 miRNAs (ex; hsa-mir-1908-5p), FKBP5 was targeted by 88 miRNAs (ex; hsa-mir-3654), CDK2 was targeted by 78 miRNAs (ex; hsa-mir-1296-5p), PLSCR1was targeted by 75 miRNAs (ex; hsa-mir-1304-5p), YAP1 was targeted by 56 miRNAs (ex; hsa-mir-548d-5p) and CDK1 was targeted by 52 miRNAs (ex; hsa-mir-103a-3p) (Table 5).

**Fig. 5.**
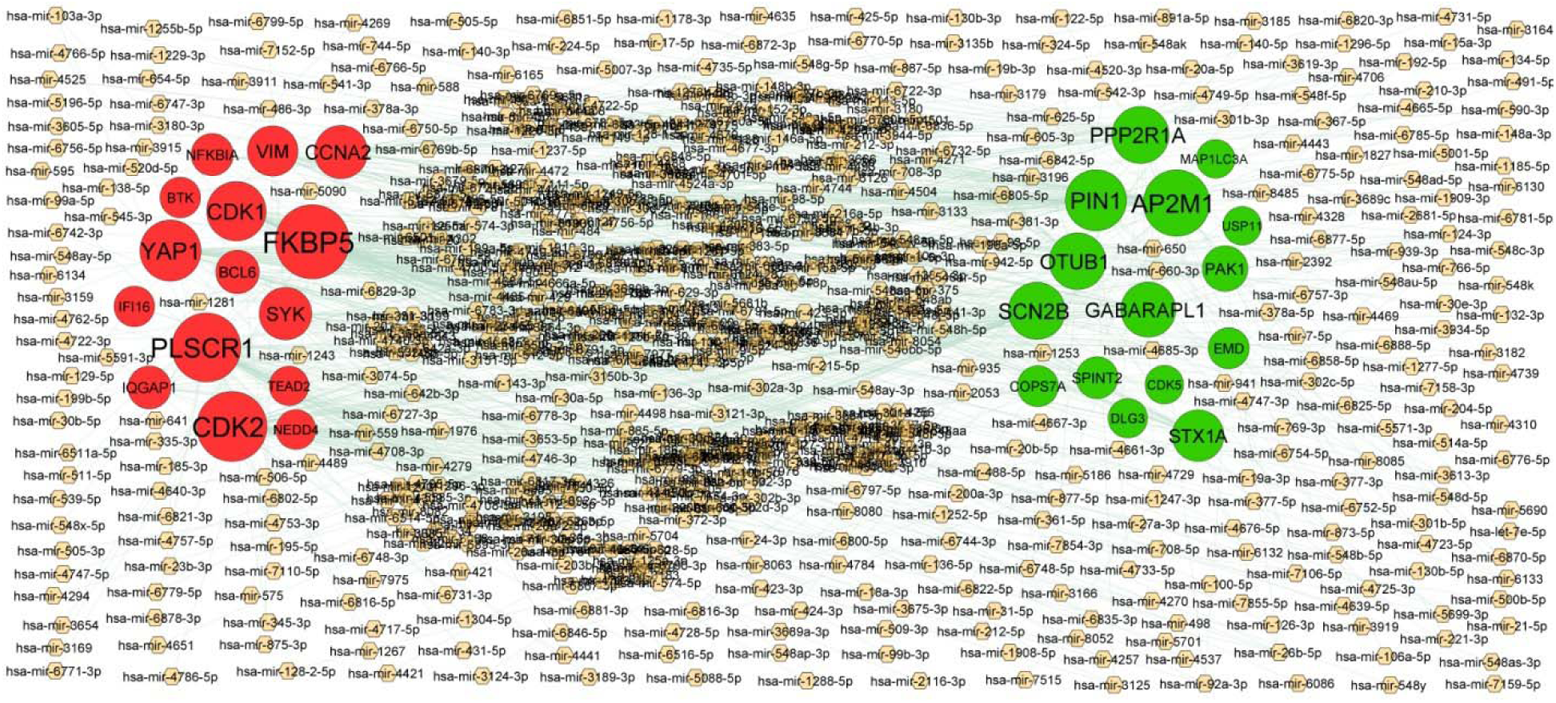
Target gene - miRNA regulatory network between target genes. The orange color diamond nodes represent the key miRNAs; up regulated genes are marked in green; down regulated genes are marked in red.

**Table 5.**
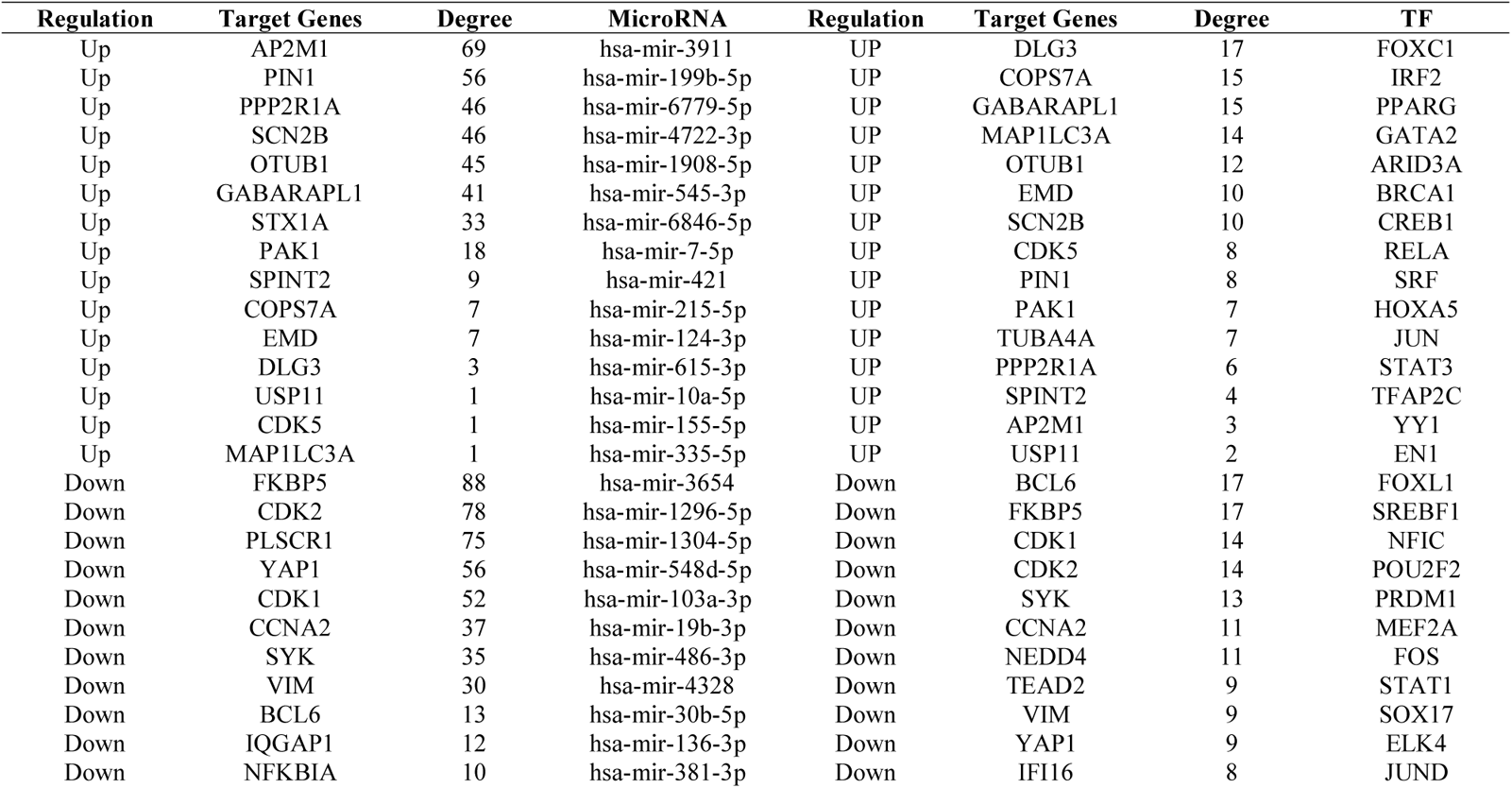
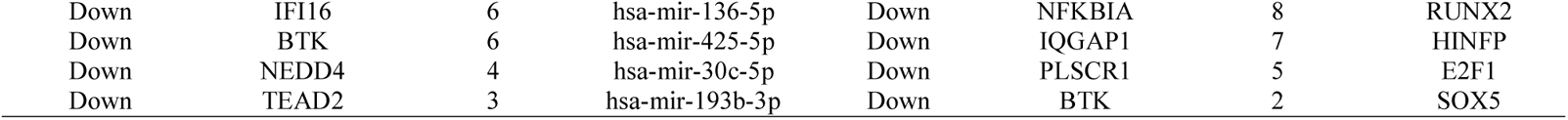
miRNA - target gene and TF - target gene interaction

### TF-hub gene regulatory network construction

TF-hub gene regulatory network was built by NetworkAnalyst for hub genes. The TF-hub gene regulatory network of hub genes was constructed with 372 (TF: 85; hub gene: 287) nodes and 2217 edges (Fig. 6). DLG3 was targeted by 17 TFs (ex; FOXC1), COPS7A was targeted by 15 TFs (ex; IRF2), GABARAPL1 was targeted by 15 TFs (ex; PPARG), MAP1LC3A was targeted by 14 TFs (ex; GATA2), OTUB1 was targeted by 12 TFs (ex; ARID3A), BCL6 was targeted by 17 TFs (ex; FOXL1), FKBP5 was targeted by 17 TFs (ex; SREBF1), CDK1 was targeted by 14 TFs (ex; NFIC), CDK2 was targeted by 14 TFs (ex; POU2F2) and SYK was targeted by 13 TFs (ex; PRDM1) (Table 5).

**Fig. 6.**
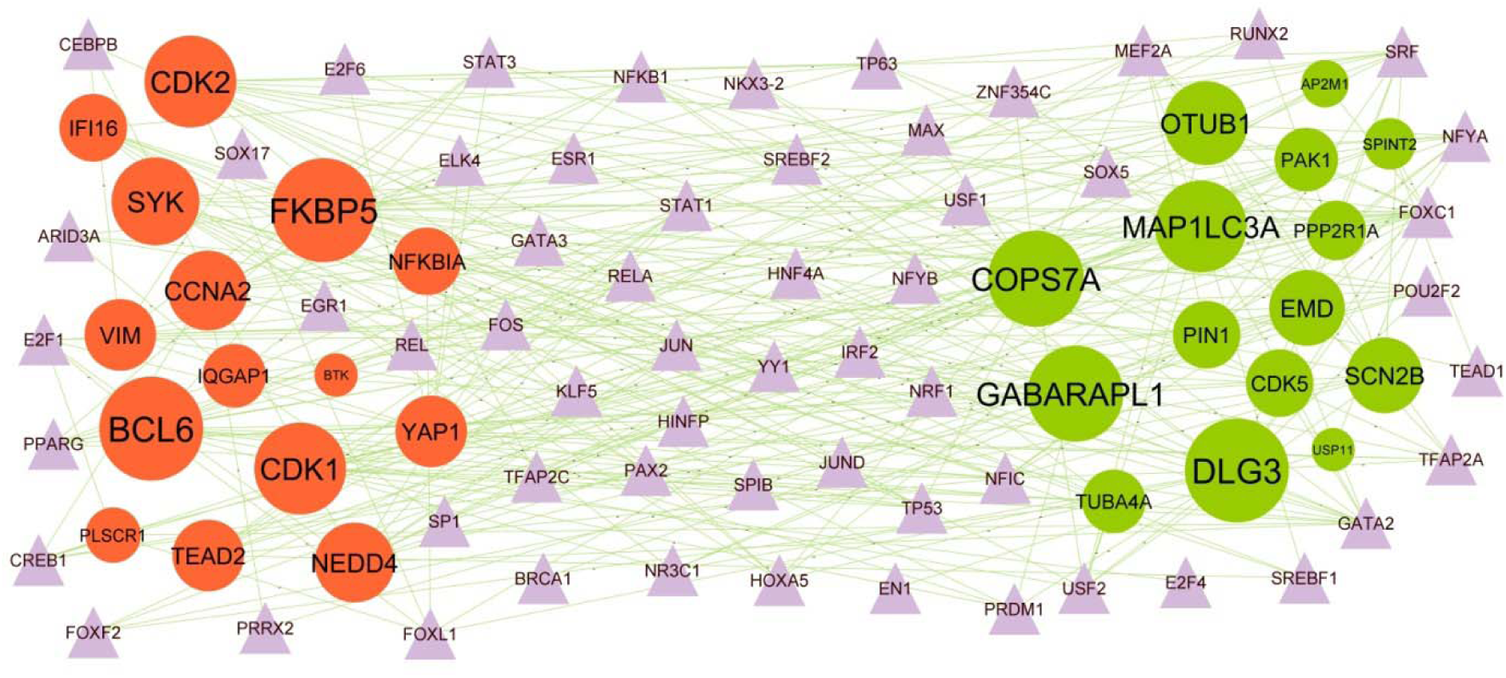
Target gene - TF regulatory network between target genes. The purple color triangle nodes represent the key TFs; up regulated genes are marked in green; down regulated genes are marked in red.

### Receiver operating characteristic curve (ROC) analysis

A ROC curve was plotted to evaluate the diagnostic value of OTUB1, PPP2R1A, AP2M1, PIN1, USP11, CDK2, IQGAP1, NEDD4, VIM and CDK1 (Fig. 7). The AUCs for the 10 hub genes were 0.943, 0.934, 0.874, 0.846, 0.854, 0.931, 0.929, 0.869, 0.940 and 0.860, respectively (Fig.7). This analysis demonstrated that the 10 hub genes had a diagnostic role in PD.

**Fig. 7.**
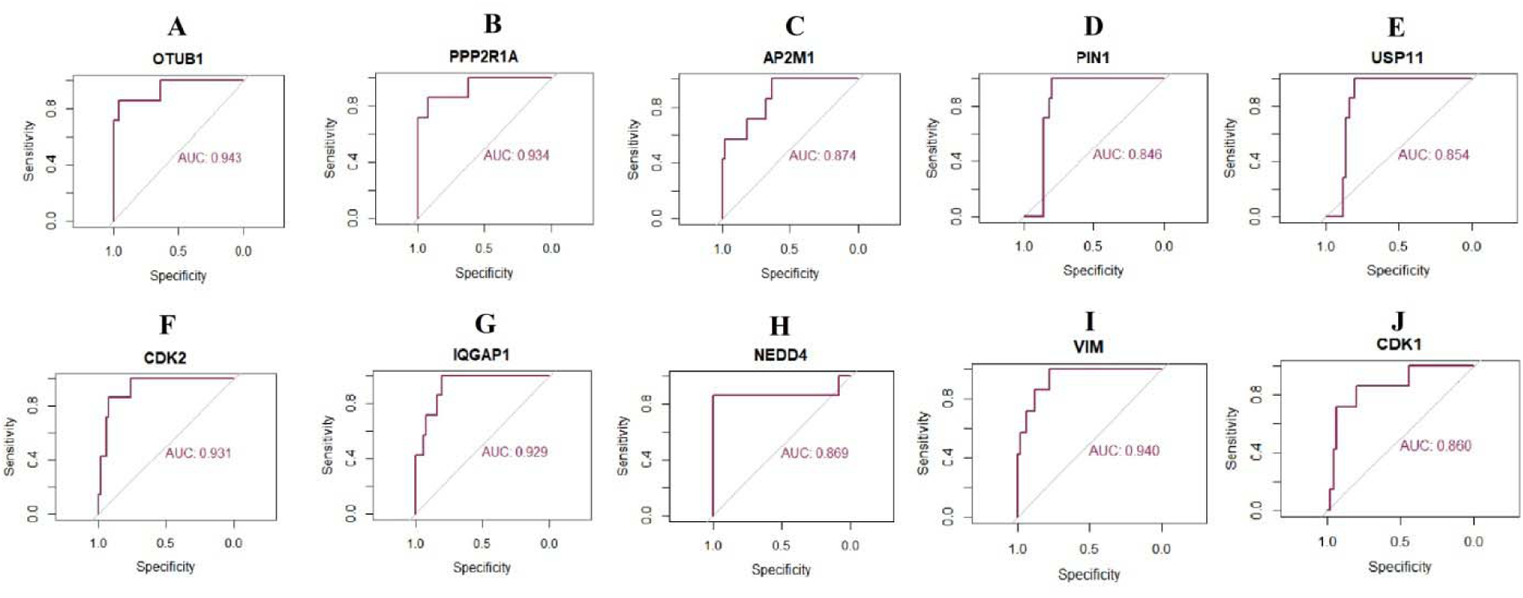
ROC curve analyses of hub genes. A) OTUB1 B) PPP2R1A C) AP2M1 D) PIN1 E) USP11 F) CDK2 G) IQGAP1 H) NEDD4 I) VIM J) CDK1

## Discussion

PD is a neurodegenerative disease characterized by tremor and bradykinesia. However, the exact mechanisms linked with PD are not clear. It has been demonstrated that tic factors play key roles in the advancement of PD. However, owing to lack of validated genetic targets, no possible therapeutic agents have been reported for the effective and safe treatment of the disease, so our goal was to find the key DEGs, associated pathways, and models that might be used as potential novel biomarkers or therapeutic targets for PD.

In the present investigation, integrated bioinformatics analysis of NGS data (GSE135036) was used to find the potential key genes related to PD. By performing DEGs analysis, 478 up regulated and 479 down regulated genes were successfully identified. Previous studies have demonstrated that VGF (VGF nerve growth factor inducible) [38] and SST (somatostatin) [39] are linked with the development mechanisms of Huntington’s disease. VGF (VGF nerve growth factor inducible) [40] and RRM2 [41] expression has significant diagnosis value in amyotrophic lateral sclerosis patients and acts as potential targets for amyotrophic lateral sclerosis targeted therapy. The recent studies have reported the identification of VGF (VGF nerve growth factor inducible) [42], CRH (corticotropin releasing hormone) [43], NPAS4 [44] and SST (somatostatin) [45] biomarkers in schizophrenia. Altered CRH (corticotropin releasing hormone) [46] and SST (somatostatin) [47] expression levels are associated with PD and are considered to be a biomarker and therapeutic target for PD. CRH (corticotropin releasing hormone) [48] and SST (somatostatin) [49] were shown to participate in facilitating Alzheimer’s disease. Kümpfel et al. [50] and Basivireddy et al. [51] found that biomarkers, including CRH (corticotropin releasing hormone) and SST (somatostatin) positively correlate with multiple sclerosis. CRH (corticotropin releasing hormone) [52] and SST (somatostatin) [53] have been reported to be altered expression in autism spectrum disorder. DUSP4 [54] have been reported to be related to epilepsy. These findings suggested that these genes might participate in the occurrence and development of PD.

In GO function and REACTOME pathway annotation, some genes involved with regulation of neurological and immune system processes were enriched in PD samples. Signaling by NTRK1 (TRKA) [55], cardiac conduction [56], signaling by GPCR [57], immune system [58], cytokine signaling in immune system [59], interferon signaling [60] and toll-like receptor cascades [61] were responsible for development of PD. Recent studies have shown that EGR2 [62], WNT1 [63], ARC (activity regulated cytoskeleton associated protein) [64], CHRNA7 [65], SEZ6L2 [66], IL1RAPL2 [67], PER2 [68], PCDH19 [69], CNTNAP2 [70], SLC12A5 [71], CDK5 [72], ACTL6B [73], GABRD (gamma-aminobutyric acid type A receptor subunit delta) [74], CACNA1G [75], HTR2C [76], STX1A [77], ATP1A3 [78], RIMS3 [79], CNTNAP2 [80], CDH8 [81], SCAMP5 [82], SYNGR1 [83], ARHGEF9 [84], DLG3 [85], RBP4 [86], IL9 [87], S100A9 [88], HGF (hepatocyte growth factor) [89], C3 [90], FKBP5 [91], GABRE (gamma-aminobutyric acid type A receptor subunit epsilon) [92], NCKAP1L [93], PIK3CG [94], ITGB3 [95], ANXA1 [96], SYNE2 [97] and DBI (diazepam binding inhibitor, acyl-CoA binding protein) [98] were closely involved with the occurrence, development, and prognosis of autism spectrum disorder. EGR2 [99], ADCYAP1 [100], CHRNA7 [101], NRN1 [102], ETV5 [103], STXBP1 [104], CAMKK2 [105], VAMP2 [106], SYNGR1 [107], NOD2 [108], TLR2 [109], BRCA2 [110] and LEF1 [111] were previously reported to be critical for the development of bipolar disorder. Accumulating evidence shows that EGR2 [112], WNT1 [113], ARC (activity regulated cytoskeleton associated protein) [114], ADCYAP1 [115], SCN5A [116], RTN4R [117], CHRNA7 [118], NRGN (neurogranin) [119], CHRM1 [120], CCK (cholecystokinin) [121], RGS4 [122], LINGO1 [123], PAK1 [124], PCDH19 [125], NRN1 [126], CX3CL1 [127], CNTNAP2 [128], SLC12A5 [129], GAS7 [130], NTNG1 [131], RAB3A [132], STXBP1 [104], CHRNB2 [133], CDK5 [134], HTR5A [135], SLC30A3 [136], HTR3B [137], HTR2C [138], TAMALIN (trafficking regulator and scaffold protein tamalin) [139], STX1A [140], GRM2 [141], SLC1A6 [142], NPTX2 [143], CAMKK2 [144], SYP (synaptophysin) [145], VAMP2 [146], ATP1A3 [147], SV2A [148], CNTNAP2 [149], CAP2[150], SYNGR1 [151], SNCB (synuclein beta) [152], RBP4 [153], KIF17 [154], CHI3L1 [155], CCR5 [156], C1QB [157], TLR7 [158], TLR2 [159], MNDA (myeloid cell nuclear differentiation antigen) [160], C3 [161], IL2RG [162], MICB (MHC class I polypeptide-related sequence B) [163], FKBP5 [164], NEFH (neurofilament heavy chain) [165], CELSR1 [166], APBB1IP [167], CD34 [168], BRCA2 [110], ITGB3 [169], ANXA3 [170], NQO1 [171], B2M [172], SLC39A12 [173], NEDD4 [174], COX2 [175], CFH (complement factor H) [176], TGFBR2 [177], MYD88 [178], ITGA8 [179], REST (RE1 silencing transcription factor) [180] and KCNJ10 [181] are altered expressed in schizophrenia. WNT1 [182], NRGN (neurogranin) [183], CCK (cholecystokinin) [184], RGS4 [185], PLK2 [186], LINGO1 [187], UNC5D [188], MEF2D [189], CX3CL1 [190], PIN1 [191], RET (ret proto-oncogene) [192], NME1 [193], STX1B [194], CDK5 [195], NPTX2 [196], VAMP2 [197], PRKAR1B [198], CAP2 [150], SNCB (synuclein beta) [199], AP2M1 [200], S100A9 [201], TLR8 [202], SERPINA1 [203], CCR5 [204], NOD2 [205], TLR7 [202], HGF (hepatocyte growth factor) [206], TLR2 [207], PTPRC (protein tyrosine phosphatase receptor type C) [208], C3 [209], LAMP3 [210], GLI1 [211], GPR4 [212], TLR1 [213], OSMR (oncostatin M receptor) [214], NFATC2 [215], GPNMB (glycoprotein nmb) [216], NQO1 [217], B2M [218], TRDN (triadin) [219], HK2 [220], NEDD4 [221], ATP6 [222], COX2 [223], CASP6 [224], MYD88 [225], NFKBIA (NFKB inhibitor alpha) [226], IL13RA1 [227], ND1 [228], TP53INP1 [229], CSF1 [230], ITPKB (inositol-trisphosphate 3-kinase B) [231], ANXA1 [232], SUMO4 [233], ITGA8 [234] and REST (RE1 silencing transcription factor) [235] have been shown to be activated in **PD**. WNT1 [236], RTN4R [237], MEF2D [238], CX3CL1 [239], PIN1 [240], UNC13A [241], CDK5 [242], SLC30A3 [243], TUBA4A [244], BCL2A1 [245], CHI3L1 [246], SERPINA1 [247], CCR5 [248], C7 [249], S100A4 [250], C1QB [251], SPP1 [252], TLR7 [253], TLR2 [254], NEFH (neurofilament heavy chain) [255], GPNMB (glycoprotein nmb) [256], B2M [257], COX2 [258], YAP1 [259], MYD88 [260], CSF1 [261], REST (RE1 silencing transcription factor) [262], DDX58 [263], LRP4 [264] and KCNJ10 [265] contributes to the progression of amyotrophic lateral sclerosis. Previous studies had shown that the altered expression of ADCYAP1 [266], CCK (cholecystokinin) [267], LINGO1 [268], CX3CL1 [269], NECTIN1 [270], IL9 [271], TLR8 [272], CCR5 [273], NOD2 [274], C7 [275], TLR7 [276], HGF (hepatocyte growth factor) [277], TLR2 [278], PTPRC (protein tyrosine phosphatase receptor type C) [279], C3 [280], IFI16 [281], GLI1 [282], CYBB (cytochrome b-245 beta chain) [283], TLR1 [284], NEFH (neurofilament heavy chain) [285], CLIC1 [286], PDK4 [287], NFATC2 [288], GPNMB (glycoprotein nmb) [289], CD58 [290], NQO1 [291], B2M [292], ANXA2 [293], FLT1 [294], IFIH1 [295], COX2 [296], NLRC5 [297], CFH (complement factor H) [298], YAP1 [299], MYD88 [300], IQGAP1 [301], ANXA1 [302] and DDX58 [303] were closely related to the occurrence of multiple sclerosis. Studies had shown that NEUROD6 [304], CHRNA7 [305], NRGN (neurogranin) [306], CCK (cholecystokinin) [307], RGS4 [308], SEZ6 [309], PLK2 [310], LINGO1 [311], NRN1 [312], CX3CL1 [313], CNTNAP2 [314], CALM3 [315], PIN1 [316], RAB3A [317], CHRNB2 [318], CDK5 [319], RPH3A [320], NPTX2 [321], NPTXR (neuronal pentraxin receptor) [322], SEZ6 [323], CAMKK2 [324], SYP (synaptophysin) [325], SV2A [326], PRKAR1B [198], CDH13 [327], CNTNAP2 [328], CALM3 [329], CAP2 [150], SLC10A4 [330], RBP4 [331], HPX (hemopexin) [332], CALHM1 [333], GNG13 [334], CHI3L1 [335], STC1 [336], FPR2 [337], S100A9 [338], CCR5 [339], C7 [340], CDK1 [341], HGF (hepatocyte growth factor) [342], TLR5 [343], TFPI (tissue factor pathway inhibitor) [344], TLR2 [345], C3 [346], CFI (complement factor I) [347], ALOX5AP [348], SELL (selectin L) [349], FKBP5 [350], CASP4 [351], SYK (spleen associated tyrosine kinase) [352], CGAS (cyclic GMP-AMP synthase) [353] NCKAP1L [354], CLIC1 [286], NFATC2 [355], CD34 [356], GPNMB (glycoprotein nmb) [357], CDK2 [358], TNFSF10 [359], BTK (Bruton tyrosine [362], MSTN (myostatin) [363], kinase) [360], NQO1 [361], CTSS (cathepsin S) [] IFITM3 [364], DOCK2 [365], BCL6 [366], COX2 [367], CASP7 [368], CFH (complement factor H) [369], YAP1 [370], TGFBR2 [371], CASP6 [372], MYD88 [373], CYTB (cytochrome b) [374], RGCC (regulator of cell cycle) [375], CSF1 [376], ITPKB (inositol-trisphosphate 3-kinase B) [377], CD2AP [378], REST (RE1 silencing transcription factor) [379] and BACE2 [380] were altered expressed in patients with Alzheimer’s disease. Byrne et al. [381], Lee et al. [382], Hays et al. [383], Rudinskiy et al. [384], Subbarayan et al. [385], Carnemolla et al. [386], Cherubini et al. [387], Wang et al. [388], Goto et al. [389], Bertoglio et al. [390], Griffioen et al. [391], Larkin and Muchowski [392], Bailus et al. [393], Sharma et al. [394], Bondulich et al. [395], Wong et al. [396], Orozco-Díaz et al. [397] and Picó et al. [398] revealed that NRGN (neurogranin), CHRM1, CCK (cholecystokinin), HPCA (hippocalcin), CX3CL1, PIN1, CDK5, GRM2, SYP (synaptophysin, SV2A, TLR2, C3, FKBP5, CGAS (cyclic GMP-AMP synthase), MSTN (myostatin), CASP6, REST (RE1 silencing transcription factor) and SLC19A3 are associated with Huntington’s disease. The expression and prognosis of TUBB2A [399], PAK1 [400], PRKCG (protein kinase C gamma) [401], CACNA1G [402], ATP1A3 [403] and KCNJ10 [404] have been investigated in ataxia. SCN3B [405], PCDH19 [406], CNTNAP2 [407], STXBP1 [408], CHRNB2 [409], NAPA (NSF attachment protein alpha) [410], STX1B [411], GABRG2 [412], CAMKK2 [144], CNTNAP2 [413], ARHGEF9 [414], COX8A [415], CALHM1 [416], SLC45A1 [417], TLR5 [418] and KCNJ10 [419] could be acuseful prognostic biomarker in epilepsy. The altered expression of NPTX2 [420], VAMP2 [421], PRKAR1B [422], SNCB (synuclein beta) [199], AP2M1 [423], TUBA4A [424], KIF17 [425] and SYK (spleen associated tyrosine kinase) [426] might be related to the progression of dementia. The above evidence revealed that these genes were related with disorders of the nervous system and might have a function in PD.

The PPI network and modules of DEGs was analyzed by HIPPIE interactome. After screening hub genes, these key genes related to PD prognosis, diagnosis amd novel therapy were identified. Reports describe the role of OTUB1 in PD [427]. Miron et al. [428] demonstrates that PPP2R1A is up-regulated in Alzheimer’s disease. However, the role of USP11, VIM, WWTR1, RASSF8 and TEAD2 in the development of PD remains unclear. Further investigations will be required to identify the relationship between these genes and PD.

miRNA-hub gene regulatory network and TF-hub gene regulatory network containing the hub genes were constructed. After screening miRNA and TFs, these key miRNA and TFs related to PD prognosis, diagnosis amd novel therapies were identified. A previous study demonstrated that hsa-mir-103a-3p [429] was altered expressed in multiple sclerosis. Wu et al [430] reported that hsa-mir-103a-3p expression migt be regarded as an indicator of susceptibility to autism spectrum disorder. Kurzawski et al [431] and Lou et al [432] demonstrated that the altered expression of GATA2 and SREBF1 are associated with prognosis in patients with PD. Study have reported that patients with SREBF1 [433] expression tended to suffer from amyotrophic lateral sclerosis. SREBF1 [434] was elevated in patients with schizophrenia. These results indicated that SCN2B, PLSCR1, COPS7A, GABARAPL1, hsa-mir-3911, hsa-mir-199b-5p, hsa-mir-6779-5p, hsa-mir-4722-3p, hsa-mir-1908-5p, hsa-mir-3654, hsa-mir-1296-5p, hsa-mir-1304-5p, hsa-mir-548d-5p, FOXC1, IRF2, PPARG, ARID3A, FOXL1, NFIC (nuclear factor 1 C), POU2F2 and PRDM1 might be a potential biomarker of PD.

In conclusion, we used a series of bioinformatics analysis methods to identify the essential genes and pathways involved in PD initiation and progression from NGS containing normal control samples and PD samples. Our results provide a more detailed molecular mechanism for the advancement of PD, shedding light on the potential biomarkers and therapeutic targets. However, the interacting mechanism and function of genes need to be confirmed in further experiments.

## Acknowledgement

I thank Peipei Li, Labrie lab, Center for Neurodegenerative Science, Van Andel Research Institute, Grand Rapids, USA, very much, the author who deposited their NGS dataset GSE135036, into the public GEO database.

## Ethical approval

This article does not contain any studies with human participants or animals performed by any of the authors.

## Informed consent

No informed consent because this study does not contain human or animals participants.

## Availability of data and materials

The datasets supporting the conclusions of this article are available in the GEO (Gene Expression Omnibus) (https://www.ncbi.nlm.nih.gov/geo/) repository. [(GSE135036) https://www.ncbi.nlm.nih.gov/geo/query/acc.cgi?acc=GSE135036)]

## Consent for publication

Not applicable.

## Author Contributions

1. B. V. - Writing original draft, and review and editing
2. C. V. - Software and investigation

